# Social learning strategies regulate the wisdom and madness of interactive crowds

**DOI:** 10.1101/326637

**Authors:** Wataru Toyokawa, Andrew Whalen, Kevin N. Laland

## Abstract

Why groups of individuals sometimes exhibit collective ‘wisdom’ and other times maladaptive ‘herding’ is an enduring conundrum. Here we show that this apparent conflict is regulated by the social learning strategies deployed. We examined the patterns of human social learning through an interactive online experiment with 699 participants, varying both task uncertainty and group size, then used hierarchical Bayesian model-ftting to identify the individual learning strategies exhibited by participants. Challenging tasks elicit greater conformity amongst individuals, with rates of copying increasing with group size, leading to high probabilities of herding amongst large groups confronted with uncertainty. Conversely, the reduced social learning of small groups, and the greater probability that social information would be accurate for less-challenging tasks, generated ‘wisdom of the crowd’ effects in other circumstances. Our model-based approach provides evidence that the likelihood of collective intelligence versus herding can be predicted, resolving a longstanding puzzle in the literature.

---

Understanding the mechanisms that account for accurate collective decision-making amongst groups of animals – ‘collective intelligence’ – has been a central focus of animal behaviour research ^1–5^. There are a large number of biological examples showing that collectives of poorly informed individuals can achieve a high performance in solving cognitive problems under uncertainty ^6–10^. Although these findings suggest fundamental cognitive benefits of grouping ^11^, there is also a long-standing recognition, especially for humans, that interacting individuals may sometimes be overwhelmed by the ‘*extraordinary popular delusions and madness of crowds*’ ^12^. Herd behaviour (i.e. an alignment of thoughts or behaviours of individuals in a group) occurs because individuals imitate each other ^13–15^, even if each is a rational decision-maker ^16^. Imitation is thought to be a cause of financial bubbles ^12;17^, ‘groupthink’ ^18^ and volatility in cultural markets ^19;20^. More generally, interdependence between individual decisions may undermine the wisdom of crowds effect ^21^ (but see ^22^), whilst potential disadvantages of information transfer are well-recognised in the biological literature ^23;24^. It seems that information transmission among individuals, and making decisions collectively, is a double-edged sword: combining decisions may provide the benefits of collecitve intelligence, but at the same time, increase the risk of an informational cascade ^16^. Collectively, an understanding of whether and, if so, how it is possible to prevent or reduce the risk of maladaptive herding, while concurrently keeping or enhancing collective intelligence, is largely lacking.

A balance between using individual and social information may play a key role in determining the trade-off between collective wisdom and ‘madness’ ^25^. If individuals are too reliant on copying others’ behaviour, any idea, even a maladaptive one, can propagate in the social group through positive feedbacks ^2;26^. For instance, disproportionally strong positive responses to recruitment signals in social insects have been shown to trap the whole colony to exploit a suboptimal, out-dated resource ^24;27^. Likewise, conformity-biased transmission in humans and other animals can potentially lead groups to converge on a maladaptive behaviour ^16;23;28;29^. On the other hand, however, if individuals completely ignore social information so as to be independent, they will fail to exploit the benefits of aggregating information through social interactions. The extent to which individuals should use social information should fall between these two extremes. Evolutionary models predict that the balance between independence and interdependence in collective decision-making may be changeable, contingent upon the individual-level flexibility and inter-individual variability associated with the social learning strategies deployed in diverse environmental states ^28;30;31^.

Experimental studies report that animals (including humans) increase their use of social information as the returns from asocial learning become more unreliable ^32–37^, whilst theory and data suggest that the benefits to individuals of social learning increase with group size ^34;38–42^. Selectivity in the predicted use of social information may impact on collective decision-making because slight differences in the parameter values of social information use are known to be able to alter qualitatively the collective behavioural dynamics ^1;2;5;43;44^. Therefore, researchers should expect populations to exhibit a higher risk of being trapped with maladaptive behaviour with increasing group size and decreasing reliability of asocial learning (and concomitant increased reliance on social learning).

From the viewpoint of the classic wisdom of crowds theory, increasing group size may increase collective accuracy ^45–48^. The relative advantage of the collective over solitary individuals may also be highlighted by increased task difficulty, because there would be more room for the performance of difficult tasks to be improved compared to easier tasks in which high accuracy can be achieved by asocial learning only. To understand the collective decision performance of social learners fully requires fine-grained quantitative studies of social learning strategies and their relations to collective dynamics, linked to sophisticated computational analysis.

The aims of this study were twofold. First, we set to test the hypothesis that the circumstances under which collective decision making will generate ‘wisdom’ can be predicted with knowledge of the precise learning strategies individuals deploy, through a combination of experimentation and theoretical modelling. The choice of an abstract decision-making task allowed us to implement a computational modelling approach, which has been increasingly deployed in quantitative studies of animal social learning strategies ^35;49–51^. In particular, computational modelling allowed us to conduct a parametric description of different information-gathering processes and to estimate the parameter values at an individual-level resolution. This approach allows us to characterise the complex relationship between individual-level decision, learning strategies and collective-level behavioural dynamics.

Second, we added resolution to our analyses by manipulating both task uncertainty and group size in our web-based experiments with adult human subjects, predicting that these factors would induce heavier use of social information in humans, and thereby alter the balance between collective intelligence and the risk of inflexible herding. To do this, we focused on human groups exposed to a simple gambling task called a multi-player ‘multi-armed bandit’, where both asocial and social sources of information were available ^35;51;52^. Through development of an interactive, web-based collective decision-making task, and use of hierarchical Bayesian statistical methods in fitting our computational model to the experimental data, we identify the individual-level learning strategies of participants as well as quantify variation in different learning parameters, allowing us to conduct an informed exploration of the population-level outcomes. The results provide clear evidence that the collective behavioural dynamics can be predicted with knowledge of human social learning strategies.

Below, we firstly deploy agent-based simulation to illustrate how the model parameters relating to social learning can in principle affect the collective-level behavioural dynamics. The simulation provides us with precise, quantitative predictions concerning the complex relationship between individual behaviour and group dynamics. Second, we present the findings of a multi-player web-based experiment with human participants that utilises the gambling task framework. Applying a hierarchical Bayesian statistical method, we estimated the model’s parameters for each of 699 different individuals, allowing us to (*i*) examine whether and, if so, how social information use is affected by different group size and task uncertainty, and (*ii*) whether and how social-information use affects both collective intelligence and the risk of maladaptive herding.

## 1 Results

### 1.1 The relationship between social learning and the collective behaviour

Figure 1 shows the relationship between the average decision accuracy and individual-level social information use obtained from our individual-based model simulations, highlighting the trade-off between accuracy and flexibility of collective decision-making. When the mean *conformity exponent* is small (i.e. *θ̄* = (*Σ_i_ θ_i_*)/*individuals* = 1), large groups are able to recover the decision accuracy quickly as do small groups after the location of the optimal option has been switched, whereas overall improvement by increasing group size in decision accuracy is subtle when the average *social learning weight* is also small (i.e. *σ̄* = (Σ_*i*_ Σ_*t*_ *σ_i_*_,*t*_)/(*individuals* × *rounds*) = 0.3; Figure 1A and 1C). On the other hand, when both the conformity exponent *θ̄* and the social learning weight *σ̄* are large, average performance is no longer monotonically improving with increasing group size, and it is under these circumstances that the strong herding effect becomes prominent (Figure 1D). Although the high conformity bias with low social learning weight makes large groups more accurate before the environment changes, larger groups are less flexible in performance recovery (Figure 1C). The patten is robust for other parameter regions (Supplementary Supplementary Figure 2).

**Figure 1:**
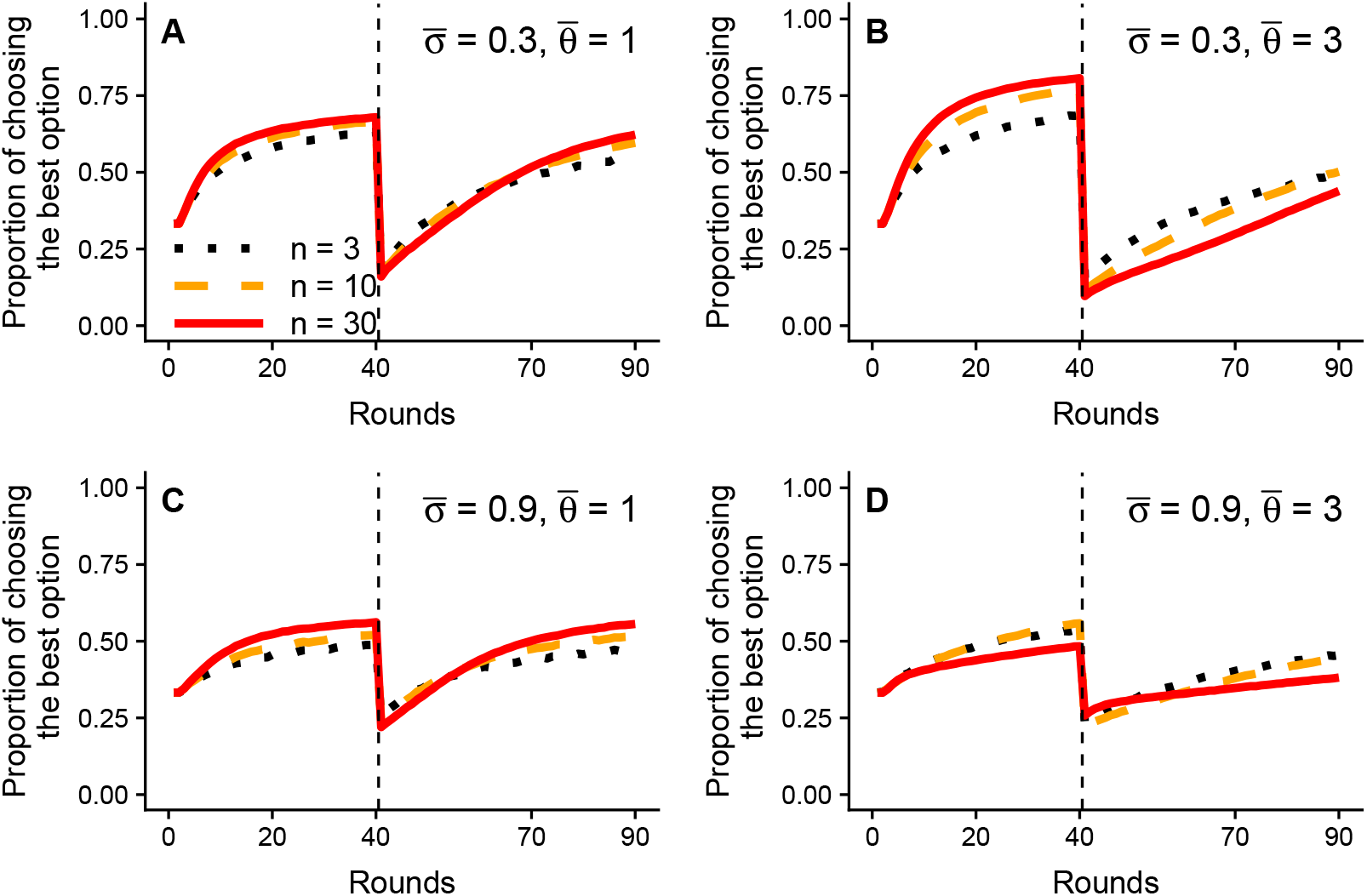
Findings of the individual-based model showing the effects of social information use on the average decision accuracy over replications. The x-axis gives the round and y-axis gives the proportion of individuals expected to choose the optimal slot (i.e. decision accuracy) averaged over all replications. The vertical dashed line indicates the timing of environmental (i.e. payoff) change (at *t* = 41). Different group sizes are shown by different styles (black (dotted): *n* = 3, orange (dashed): *n* = 10, red (solid): *n* = 30). We set the average slopes for the *social learning weight* to be equal to zero for the sake of simplicity; namely, *μ_δ_* = 0. Other free parameter values (i.e. *μ_α_*, 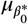, *μ_ε_ v_a_*, 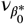, *v_ε_*, *v_σ_*, *v_δ_* and *v_θ_*) are best approximates to the experimental fitted values (see Table 2 and Supplementary table 1).

Figure 2C and 2D indicate that when both *θ̄* and *σ̄* are large the collective choices converged either on the good option or on one of the poor options almost randomly, regardless of the option’s quality, and that once individuals start converging on an option the population gets stuck. As a result, the distribution of the groups’ average performance over the replications becomes a bimodal ‘U-shape’. Interestingly, however, the maladaptive herding effect remains relatively weak in smaller groups (see Figure 1D; the dotted line). This is because the majority of individuals in smaller groups (i.e. two individuals out of three) are more likely to break the cultural inertia by simultaneously exploring another option by chance than are the majority in larger groups (e.g. six out of ten).

**Figure 2:**
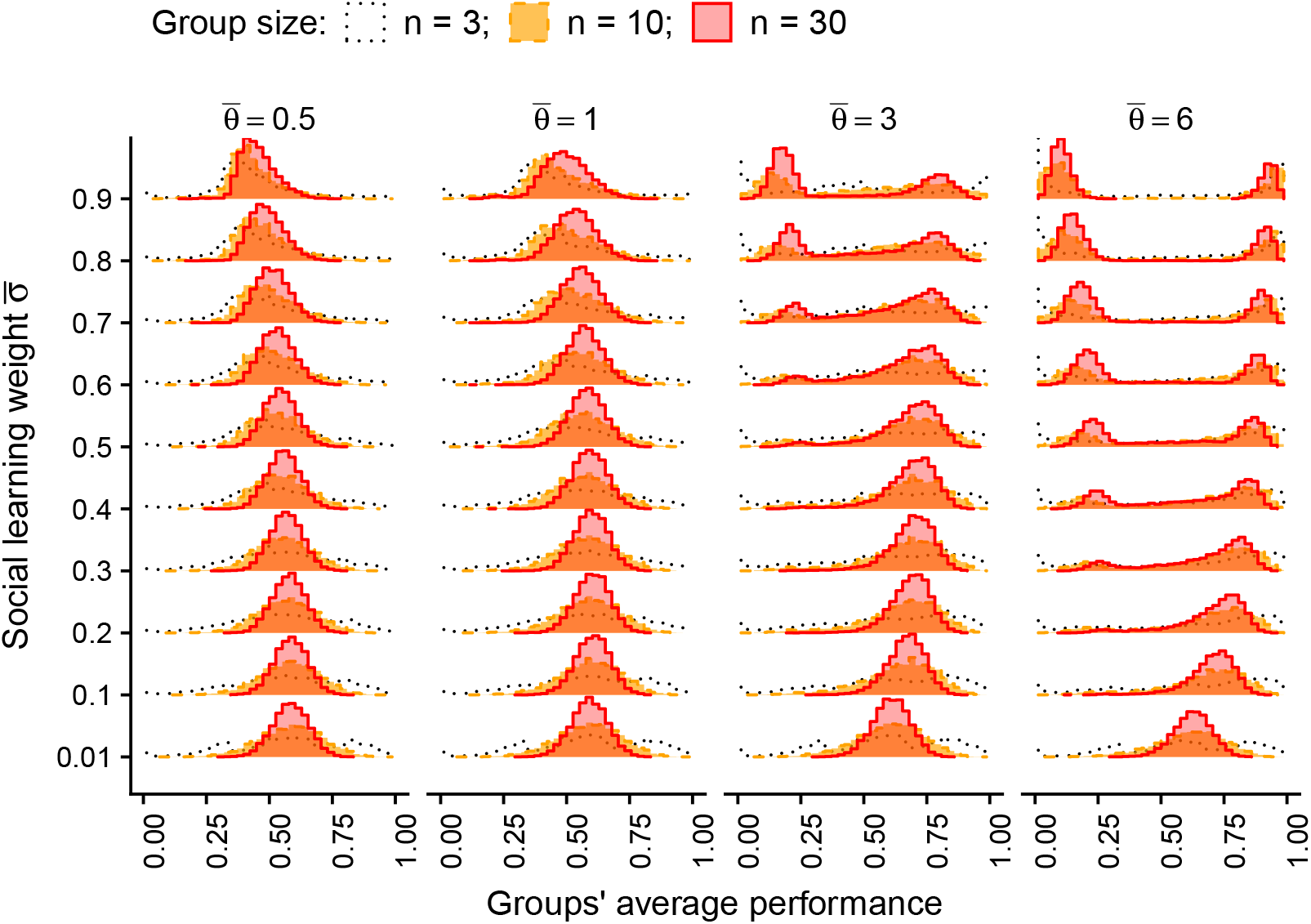
Results from the individual-based model simulations showing the distribution of each group’s mean accuracy before environmental change (*t* ≤ 40). The x-axis gives the mean decision accuracy over the first 40 rounds (i.e. the environment 1) for each replication. Different group sizes are shown by different styles (black (dotted): *n* = 3, orange (dashed): *n* = 10, red (solid): *n* = 30). The other free parameter values are the same as in Figure 1.

In summary, the model simulation suggests an interaction between social learning weight *σ̄* and conformity exponent *θ̄* on decision accuracy and the risk of inflexible herding. When the conformity exponent is not too large, increasing group size can increase decision accuracy while concurrently retaining decision flexibility across a broad range of the mean social learning weights. When the conformity bias becomes large, however, the risk of inflexible herding arises, and, when both social learning parameters are large, collective intelligence is rare and inflexible herd behaviour dominates.

### 1.2 Collective performance of human participants

Figure 3A shows behavioural dynamics of human participants in different group sizes and different task uncertainty conditions (see Supplementary Supplementary Figure 3 for each group’s behaviour). The average decision performance of collectives (i.e. group size ≥ 2) exceeded that of solitary individuals (i.e. group size = 1) in the Moderate-uncertainty condition (i.e. the 95% Bayesian CI of *ξ_t_* exceeds 0 at regions *t ∈* [9, 40] and [67, 70]; Figure 3B). In other uncertainty conditions, no global positive effect of grouping was observed, suggesting that collective intelligence was prominent only in the Moderate-uncertainty condition. However, the main effect of group size was positive in the post-change period of the Low-uncertainty condition (mean and the 95% Bayesian CI of *ω*_2_ = 0.67 [0.44, 0.91]; Table 1), suggesting that the average performance of large groups (e.g. 12 ≤ group size ≤ 16) were better, and hence more flexible, than smaller groups and solitaries (Figure 3A). On the other hand, in the Moderate-uncertainty condition, the average performance of the collectives dropped below that of the solitaries after the environmental change (i.e. *ξ_t_ <* 0 at a region *t* ∈ [42, 45]; Figure 3B). Also, the main effect of group size was negative in the post-change period (mean and the 95% Bayesian CI of *ω*_2_ = -0.26 [-0.44, -0.11]; Table 1), suggesting that larger groups were more likely to get stuck in the out-dated option in the Moderate-uncertainty condition. In the High-uncertainty condition, the main effect of group size was positive in the prior-change period and negative post-change (mean and the 95% Bayesian CIs are *ω*_1_ = 0.07 [0.00, 0.15] and *ω*_2_ = -0.10 [-0.17, -0.02]; Table 1), although the effect size was too small to differentiate performances of different group sizes visually (Figure 3A). Using monetary earnings as an outcome variable of decision performance did not change our conclusions qualitatively (supporting Supplementary Figure 4 and Supplementary table 2).

**Table 1:** The mean and the 95% Bayesian credible intervals of the posterior for the group size effect at the phenomenological logistic model

|  | Low Uncertainty |  | Moderate Uncertainty |  | High Uncertainty |  |
| --- | --- | --- | --- | --- | --- | --- |
| $\omega_1$ | 0.08 | [-0.15, 0.33] | 0.10 | [-0.06, 0.26] | 0.07 | [0.00, 0.15] |
| $\omega_2$ | 0.67 | [0.44, 0.91] | -0.26 | [-0.44, -0.11] | -0.10 | [-0.17, -0.02] |

Note: All $\hat{R}$ values are 1.0 and the effective sample sizes are larger than 837.

**Figure 3:**
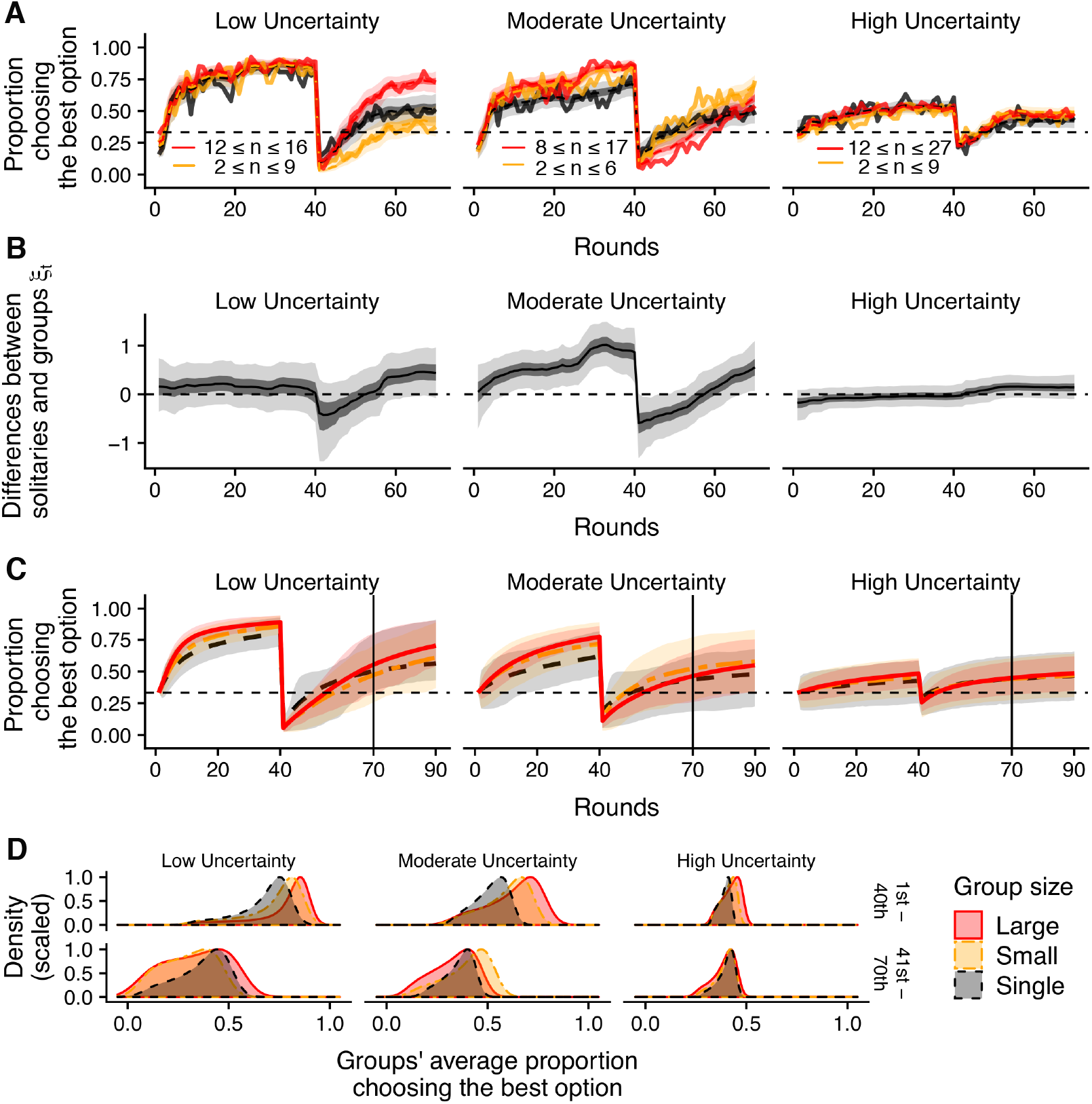
Time evolutions and distributions of decision performance for each condition. **A:** The average decision accuracies of the experimental participants (red: large groups, orange: small groups, dark grey: lone individuals). All individual performances were averaged within the same size category (solid lines). The light-shaded areas, dark-shaded areas, and dashed curves show the 95%, 50%, and median Bayesian credible intervals of the phenomenological, time-series logistic regression. Sample sizes for large, small, and lone groups are: *N* = 43, *N* = 44 and *N* = 38 for the Low-uncertainty condition; *N* = 52, *N* = 56 and *N* = 37 for the Moderate-uncertainty condition; and *N* = 259, *N* = 168 and *N* = 58 for the High-uncertainty condition, respectively. **B:** Change in the main effect of the dummy variable of grouping on the decision accuracy at the phenomenological regression model. The shaded areas are the Bayesian CIs and solid curves are the median. **C, D:** Change and distribution in average decision accuracy of the individual-based post-hoc simulations of the learning process model using the experimentally fit parameter values. **C:** All replications were averaged within the same size category (solid lines). The shaded areas give the 50% quantiles. The experimental horizon (i.e. *t* = 70) is indicated by the vertical line. **D:** Performance was averaged within prior-and post-change periods for each replication for each group sizes category.

Our phenomenological model regression established that manipulating both task uncertainty and group size indeed affected the collective decision dynamics. Below, we address whether or not the pattern could be explained with knowledge of human social learning strategies estimated through our learning and decision-making computational model.

### 1.3 Estimation of human social information use

Using posterior estimation values obtained by the hierarchical Bayesian model fitting method (Table 2), we were able to categorise the participants as deploying one of three different learning strategies based on their fitted conformity exponent values; namely, the ‘positive frequency-dependent copying’ strategy (*θ_i_* ≫ 0), the ‘negative-frequency dependent copying’ strategy (*θ_i_* ≪ 0) and the ‘random choice’ strategy (*θ_i_* ≈ 0). Note that we could not reliably detect the ‘weak positive’ frequency-dependent strategy (0 < *θ_i_* ≤ 1) due to the limitation of statistical power (Supplementary Figure 5). Some individuals whose ‘true’ conformity exponent fell between zero and one would have been categorised as exhibiting a random choice strategy (Supplementary Figure 7). Individuals identified as exhibiting a positive frequency-dependent copiers were mainly those whose conformity exponent was larger than one (*θ_i_ >* 1).

**Table 2:** The mean and the 95% Bayesian credible intervals of the posterior global means for the parameter values. The number of participants (*N*) for each experimental condition are also shown.

|  | Groups |  |  | Solitary individuals |  |  |
| --- | --- | --- | --- | --- | --- | --- |
| Uncertainty: | Low | Moderate | High | Low | Moderate | High |
| $\mu_{\alpha^*}$ (learning rate) | 0.99<br>[0.34, 1.73] | 0.90<br>[0.43, 1.44] | 0.61<br>[0.21, 1.03] | 0.85<br>[-0.07, 1.95] | -0.17<br>[-1.27, 0.89] | 0.46<br>[-0.39, 1.36] |
| $\mu_{\beta_0^*}$ (inv. temp.) | 1.84<br>[1.15, 2.70] | 1.68<br>[1.25, 2.18] | 1.38<br>[1.16, 1.62] | 1.10<br>[0.69, 1.54] | 1.44<br>[0.80, 2.07] | 0.85<br>[0.46, 1.22] |
| $\mu_{\epsilon}$ (inv. temp.) | 3.70<br>[1.98, 5.71] | 3.01<br>[1.88, 4.27] | 2.97<br>[2.37, 3.60] | 2.39<br>[1.46, 3.53] | 2.81<br>[1.64, 4.07] | 2.27<br>[1.40, 3.31] |
| $\mu_{\sigma_0^*}$ (soc. wight) | -1.55<br>[-2.71, -0.71] | -2.37<br>[-4.12, -1.01] | -2.16<br>[-2.81, -1.63] | —<br>— | —<br>— | —<br>— |
| $\mu_{\delta}$ (soc. wight) | -1.39<br>[-2.66, -0.03] | -1.55<br>[-4.29, 0.91] | -1.87<br>[-3.04, -0.81] | —<br>— | —<br>— | —<br>— |
| $\mu_{\theta}$ (conformity coeff.) | 1.65<br>[0.83, 2.82] | 3.00<br>[1.57, 4.85] | 2.67<br>[1.80, 3.73] | —<br>— | —<br>— | —<br>— |
| $N$ | 77 | 98 | 398 | 36 | 34 | 56 |

Figure 4A show the estimated frequencies of different learning strategies. Generally speaking, participants were more likely to utilize a positive frequency-dependent copying strategy than the other two strategies (the 95% Bayesian CI of the intercept of the GLMM predicting the probability to use the positive frequency-dependent copying strategy is above zero, [1.05, 2.50]; Supplementary table 4). We found that positive frequency-dependent copying decreased with increasing task uncertainty (the 95% Bayesian CI of task uncertainty effect is below zero, [-1.88, -0.25]; Supplementary table 4). We found no clear effects of either the group size, age or gender on adoption of the positive frequency-dependent copying strategy, except for the negative interaction effect between age and task uncertainty (the 95% Bayesian CI of the age × uncertainty interaction = [-1.46, -0.15]; Supplementary table 4).

**Figure 4:**
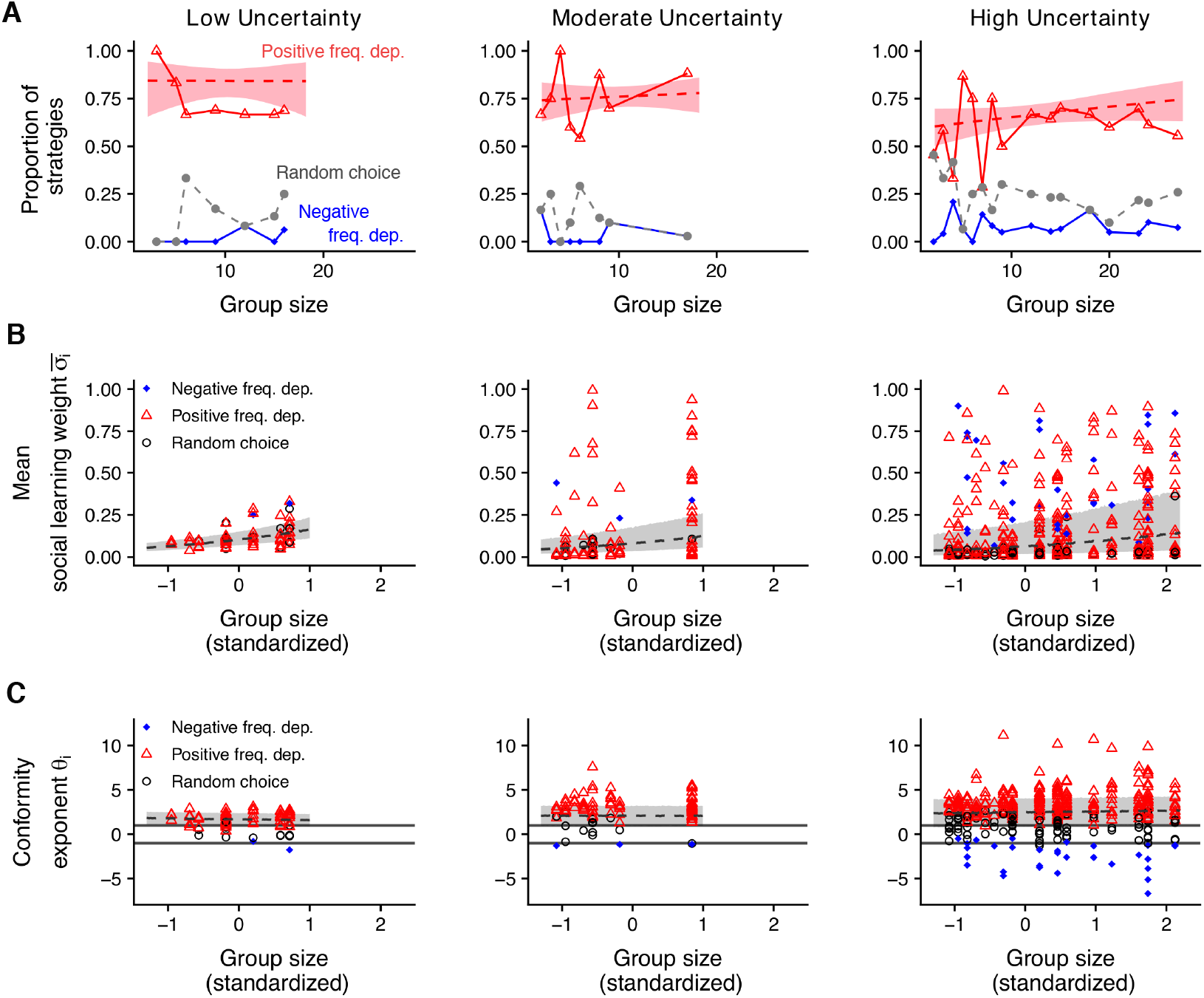
Model fitting for the three different task’s uncertain conditions (the Low-, Moderate-and High-uncertainty) and the different group size. Three different learning strategies are shown in different styles (red-triangle: positive frequency-dependent learning, blue-circle: negative frequency-dependent learning; grey-circle: nearly random choice strategy). (A) Frequencies of three different learning strategies. Note that a sum of the frequencies of these three strategies in the same group size does not necessarily equal to 1, because there are a small number of individuals eliminated from this analysis due to insufficient data. (B) Estimated social learning weight, and (C) estimated conformity exponent, for each individual shown for each learning strategy. The 50% Bayesian CIs of the fitted GLMMs are shown by dashed lines and shaded areas. The horizontal lines in (C) show a region −1 *< θ_i_ <* 1. Sample sizes for Negative Frequency Dependent, Positive Frequency Dependent, and Random Choice strategies are: *N* = 2, *N* = 61 and *N* = 14 for the Low-uncertainty condition; *N* = 3, *N* = 80 and *N* = 15 for the Moderate-uncertainty condition; and *N* = 32, *N* = 260 and *N* = 106 for the High-uncertainty condition, respectively.

We also investigated the effects of group size and task uncertainty on the fitted individual parameter values. We found that the individual mean social learning weight parameter (i.e. *σ̄_i_* = (*Σ*_t_ σ_*i*,*t*_)/(total rounds)) increased with group size (the 95% Bayesian CI = [0.15, 0.93]; Figure 4B; Supplementary table 5), and decreased with uncertainty (the 95% Bayesian CI = [-0.98, -0.22]), and age of subject (the 95% Bayesian CI = [-0.36, -0.02]). However, the negative effects of task uncertainty and age disappeared when we focused only on *σ̄_i_* of the positive frequency-dependent copying individuals, and only the positive effect of the group size was confirmed (Supplementary table 6; Supplementary Figure 6). It is worth noting that the meaning of the social learning weight is different between these three different strategies: The social learning weight regulates positive reactions to the majorities’ behaviour for positive frequency-dependent copiers, whereas it regulates avoidance of the majority for negative-frequency dependent copiers, and determines the probability of random decision-making for the random choice strategists.

The individual conformity exponent parameter *θ_i_* increased with task uncertainty (the 95% Bayesian CI = [0.38, 1.41]), but we found no significant effects of group size, age, gender or interactions (Figure 4C; Supplementary table 7). These results were qualitatively unchanged when we focused only on the positive frequency-dependent copying individuals (Supplementary table 8; Supplementary Figure 6).

We observed extensive individual variation in social information use. The greater the task’s uncertainty, the larger were individual variances in both the mean social learning weight and the conformity exponent (the 95% Bayesian CI of the GLMM’s variation parameter for *σ̄_i_* was [1.11, 1.62] (Supplementary table 5) and for *θ_i_* was [1.07, 1.54] (Supplementary table 7)). This was confirmed when focusing only on the positive frequency-dependent copying individuals: The Bayesian 95% CIs were [1.14, 1.80] (Supplementary table 6) and [0.71, 1.10] (Supplementary table 8), respectively.

The manner in which individual variation in social-information use of positive frequency-dependent copying individuals changes over time is visualised in Figure 5. The social learning weights generally decreased with experimental round. However, some individuals in the Moderate-and the High-uncertain conditions accelerated rather than decreased their reliance on social learning over time. Interestingly, those accelerating individuals tended to have a larger conformity exponent (Supplementary Figure 5). In addition, the time-dependent *θ_i_*_,*t*_in our alternative model generally increased with experimental round in the Moderate-and the High-uncertainty conditions (Supplementary Figure 10), although the fitting of *θ_i,t_* in the alternative model was relatively unreliable (Supplementary Figure 9). These findings suggest that conformists tended to use asocial learning at the outset (i.e. exploration asocially) but increasingly started to conform as the task proceeded (i.e. exploitation socially).

**Figure 5:**
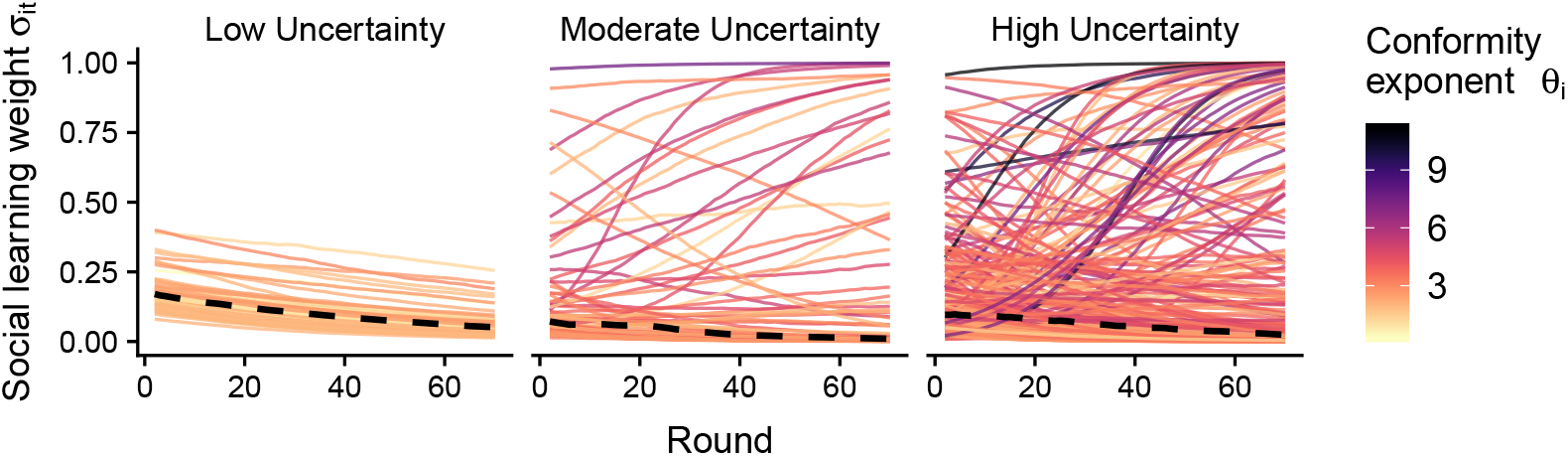
Change in fitted values (i.e. median of the Bayesian posterior distribution) of the social learning weight *σ_i_*_,*t*_ with time for each Positive Frequency Dependent individual, for each level of task uncertainty. Thick dashed lines are the median values of *σ_i_*_,*t*_across the subjects for each uncertainty condition. Individual conformity exponent values *θ_i_* are shown in different colours (higher θ_*i*_is darker). Sample size for each task uncertainty condition is: *N* = 61 (Low-uncertainty), *N* = 80 (Moderate-uncertainty) and *N* = 260 (High-uncertainty).

Extensive variation in the temporal dynamics of the social learning weight *σ_i,t_* was also found for the negative-frequency dependent copying individuals but not found for random choice individuals (Supplementary Figure 5). Individuals deploying a random choice strategy exhibited a *σ_i_*_,*t*_that approached to zero, indicating that their decision-making increasingly relied exclusively on the softmax choice rule, rather than unguided random choices, as the task proceeded.

No significant fixed effects were found in other asocial learning parameters such as the learning rate *α_i_* and the mean inverse temperature *β̄i* = (Σ*t β_i_*_,*t*_)/(total rounds) (Supplementary table 9, Supplementary table 10 and Supplementary Figure 6).

In summary, our experiments on adult humans revealed asymmetric influences of increasing task uncertainty and increasing group size on the social learning parameters. The conformity exponent increased with task uncertainty on average but the proportion of positive frequency-dependent copying individuals showed a corresponding decrease, due to the extensive individual variation emerging in the High-uncertain condition. Conversely, group size had a positive effect on the mean social learning weight, but did not affect conformity.

### 1.4 Social learning strategies explain the collective dynamics

The post-hoc simulation provides statistical predictions on how likely it is, given the fitted learning model parameters, that groups of individuals make accurate decisions and that they exhibit inflexible herding. Figure 3C shows the change over time in performance with different group sizes and different uncertainty conditions, generated by the post-hoc simulation (see also Supplementary Figure 3). The trajectories of the simulated dynamics recovered nicely the pattern observed in the experiment (Figure 3A and 3C), suggesting that the strategic changes in the individual-level social information use (Figure 4) could explain the collective-level behavioural pattern.

Figure 3D shows that larger groups are more likely to make accurate decisions than are both small groups and solitaries in the period prior to change across all uncertainty conditions, suggesting collective intelligence was operating. In the post-change period, however, performance differed between the conditions. In the Low-uncertainty condition, where we found that the participants were most likely to have a relatively weak positive frequency-dependence (i.e. *θ̄* = 1.65), large groups performed better than did small groups over 59.5% of total 10,000 repetitions. However, in the Moderate-uncertainty condition, where we found that participants were most likely to have strong positive frequency dependence (*θ̄* = 3.00, c.f. 1.65 in the Low-uncertainty condition), the large groups were more likely to get stuck on the suboptimal option, and hence the small groups performed better than did the large groups over 69.5% of repetitions (Figure 3D). The decision accuracy did not substantially differ with group size in the post-change period in the High-uncertainty condition although the large groups performed slightly better than did the small groups (50.8% of the repetitions).

Interestingly, although their relatively low conformity biases, there were some groups in the Low-uncertainty condition that seemed to exhibit herding (the ‘humped’ area at the lefthand side to the peak of the performance distribution in the post-change period; Figure 3D). This might be due to the lower softmax exploration rates among social learners in the Low-uncertainty condition (i.e. both 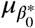 and *μ_ϵ_* were large; Table 2): the whole population gets stuck because all individuals are very exploitative on their past experience.

## 2 Discussion

We investigated whether and how human social learning strategies regulate the trade-off between collective intelligence and inflexible herding behaviour using a collective learning-and-decision-making task combined with simulation and model fitting. We examined whether manipulating the reliability of asocial learning and group size would affect the use of social information, and thereby alter the collective human decision dynamics, as suggested by our computational model simulation. Although a theoretical study has suggested that reliance on social learning and conformity bias would play a role in collective dynamics ^2;5;53^, thus far no empirical studies have quantitatively investigated the population-level consequences of these two different social learning processes. Our high-resolution, model-based behavioural analysis using a hierarchical Bayesian statistics enabled us to identify individual-level patterns and variation of different learning parameters and to explore their population-level outcomes. The results provide quantitative support for our hypothesis that the collective decision performance can be predicted with quantitative knowledge of social learning strategies.

Overall, our individual-based computational model recovered the behavioural pattern suggested by the phenomenological regression (Figure 3). Using the post-hoc simulation with individually-fit model parameters, we confirmed that in the Low-uncertainty condition, where individuals had weaker positive frequency bias (i.e. *θ̄* ≈ 1.65), larger groups were able to be more accurate than smaller groups while retaining flexibility in their decision-making ^9^, although their low asocial exploration rates seemed to undermine the potential flexibility. However, in the Moderate-and the High-uncertain conditions where individuals had the higher conformity exponent parameters (i.e. *θ̄* ≈ 3.0 and 2.7, respectively), larger groups performed better prior to environmental change but were vulnerable to getting stuck with an out-dated maladaptive option post change. Therefore, the changes in the level of conformity in human individuals ^34;41^ indeed incurred a trade-off between the collective intelligence effect and the risk of inflexible herding.

Although the social learning weight increased with increasing group size, the overall mean value was *σ̄_i_* ≈ 0.3 (Figure 4B; Supplementary Figure 5; Supplementary Figure 6) and it decreased on average as the task proceeded (Figure 5). This implies a weaker social than asocial influence on decision-making as reported in several other experimental studies ^35;54–56^ although evolutionary models tend to predict heavier reliance on social learning than experimental studies report ^57;58^. Thanks to this relatively weak reliance of social learning, the kind of extreme herding that would have blindly led a group to any option regardless of its quality, such as the ‘symmetry breaking’ known in trail-laying ant collective foraging systems ^2;5;26^, did not occur (Figure 2).

Individual differences in rates of exploration might also help to mitigate potential herding. Although a majority of participants adopted a positive frequency-dependent copying strategy, some individuals exhibited negative frequency dependence or random decision-making (Figure 4A). The random choice strategy was associated with more exploration than the other strategies, because it led to an almost random choice at a rate *σ_i_*, irrespective of the options’ quality. Negative-frequency dependent copying individuals could also be highly exploratory. These individuals tended to avoid choosing an option upon which other people had converged and would explore the other two ‘unpopular’ options. Interestingly, in the High-uncertain condition the mean social learning weights of the negative-frequency dependent copying individuals (*σ̄_i_* ≈ 0.5) were larger than that of the other two strategies (*σ̄_i_* ≈ 0.1, Supplementary Figure 5), indicating that these individuals engaged in such majority-avoiding exploration relatively frequently. Such a high variety in social information use ^59–62^and exploratory tendencies would prevent individuals from converging on a single option, leading to a mitigation of herding but concurrently diminishing the decision accuracy in high-uncertainty circumstances (Figure 3).

A methodological advantage of using computational models to study social learning strategies is its explicitness of assumptions about the temporal dynamics of behaviour, which enabled us to distinguish different learning strategies ^63–65^. For example, very exploitative asocial reinforcement learners (i.e. for whom exploitation parameter *β_i_*_,*t*_is large and the social learning weight *σ_i_*_,*t*_is nearly zero, as seen in the Low-uncertainty condition) and conformity-biased social learners (where the conformity exponent *θ_i_* is large and *σ_i_*_,*t*_is positive, as seen in the Moderate-uncertain condition) would eventually converge on the same option, resulting in the same final behavioural steady state. However, how they explored the environment, as well as how they reacted to the other individuals in the same group, are significantly different and they could produce qualitatively different collective temporal dynamics.

However, our computational model could not fully capture other, potentially more sophisticated forms of social learning strategies that participants might deploy, which might be a reason for the seemingly low rate of social learning observed in the experiment compared to theory ^57;58^. Indeed, the post-hoc simulation sometimes failed to recover the observed behavioural trajectories. In particular, experimental groups with *n* = 12, *n* = 16, and one group in *n* = 9, in the Low-uncertainty condition performed very well, exceeding the 95% CIs of the post-hoc simulation after the environmental change (Supplementary Figure 3). This indicates that collective behaviour in these groups was more flexible than our model predicted. Further empirical studies that consider a wider range of possible social learning strategies, e.g. ‘copy-rapidly-increasing-option’ strategy ^66^ or Bayesian updating ^57;67^, are needed to explore computational underpinnings of social learning and collective behaviour.

The Internet-based experimentation allowed us to conduct a real-time interactive behavioural task with larger subject pools than a conventional laboratory-based experiment. This enabled us not only to quantify the individual-level learning-and-decision processes ^68^ but also to map these individual-level processes on to the larger-scale collective behaviour ^5;15;20^. Although there are always questions about the validity of participants’ behaviour when recruited via web-based tools, we believe that the computational modelling approach coupled with higher statistical power due to the large sample size, compensates for any drawbacks. The fact that our learning model could approximate the participants’ decision trajectories effectively suggest that most of the participants engaged seriously with solving the task. An increasing body of evidence supports the argument that web-based behavioural experiments are as reliable as results from the laboratory ^69;70^.

The diverse effects of social influence on the collective wisdom of a group has been drawing substantial attention ^19;21;22;71;72^. The bulk of this literature, including many jury models and election models ^45;73^, has focused primarily on the static estimation problem, where the ‘truth’ is fixed from the outset. However, in reality, there are many situations under which the state of the true value is changing over time so that monitoring and tracking the pattern of change is a crucial determinant of decision performance ^74^. In such temporally dynamic environments, decision-making and learning are coordinated to affect future behavioural outcomes recursively ^75^. Our experimental task provides a simple vehicle for exploring collective intelligence in a dynamic situation, which encompasses this learning-and-decision-making feedback loop. Potentially, integrating the wisdom of crowds with social learning and collective dynamics research will facilitate the more tractable use of collective intelligence in a temporary changing world.

In summary, a combination of experimentation and theoretical modelling sheds new light on when groups of individuals will exhibit the wisdom of the crowds and when inflexible herding. Our analysis implies that herding is most likely amongst individuals in large groups exposed to challenging tasks. That is because challenging tasks lead to greater uncertainty and thereby elicit greater conformist learning amongst individuals, whilst rates of copying increase with group size. Difficult tasks, by definition, render identification of the optimal behavior harder, allowing groups sometimes to converge on maladaptive outcomes. Conversely, the reduced conformity levels of individuals in small groups, and the greater probability that social information would be accurate for less-challenging tasks, generated ‘wisdom of the crowd’ effects in most other circumstances. Our findings provide clear evidence that the conflict between collective intelligence and maladaptive herding can be predicted with knowledge of human social learning strategies.

## 3 Material and methods

### 3.1 Participants

The experimental procedure was approved by the Ethics Committee at the University of St Andrews (BL10808). A total of 755 subjects (354 females, 377 males, 2 others and 22 unspecified; mean age (1 *SD.*) = 34.33 (10.9)) participated through Amazon’s Mechanical Turk. All participants consented to participation through an online consent form at the beginning of the task. We excluded subjects who disconnected to the online task before completing at least the first 30 rounds from our computational-model fitting analysis due to unreliability of the model-parameter estimation, resulted in 699 subjects (573 subjects entered the group (i.e. *N* ≥ 2) and 126 entered the solitary (i.e. *N* = 1) condition). The task was only available for individuals who had greater than or equal to 90% HIT approval rate and who accessed from the United States. Although no sample-size calculation was performed in advance, our parameter recovery test confirmed that the sample size was sufficient for estimation of individual parameters using a hierarchical Bayesian method.

### 3.2 Design of the experimental manipulations

The three uncertainty conditions were: Low-uncertainty condition (differences between mean payoffs were 1.264), Moderate-uncertainty condition (differences between mean payoffs were 0.742) and High-uncertainty condition (differences between mean payoffs were 0.3). The mean payoff associated with the ‘excellent’ slot in all three conditions was fixed to 3.1 cents (Supplementary Figure 1). Each task uncertainty condition was randomly assigned for each different HIT session, and participants were allowed to participate in one HIT only. Sample size after the data exclusion for each uncertainty condition was: *N* = 113 (Low-uncertainty condition), *N* = 132 (Moderate-Uncertain condition), and *N* = 454 (High-uncertain condition). We assigned more sessions to the High-uncertainty condition compared to the other two because we expected that larger group sizes would be needed to generate the collective wisdom in noisier environments.

To manipulate the size of each group, we varied the capacity of the waiting room from 10 to 30. Because the task was being advertised on the Worker website at AMT for approximately 2 hours, some participants occasionally arrived after the earlier groups had already started. In that case the participant entered the newly opened waiting room which was open for the next 5 minutes. The number of participants arriving declined with time because newly posted alternative HITs were advertised on the top of the task list, which decreased our task’s visibility. This meant that a later-starting session tended to begin before reaching maximum room capacity, resulting in the smaller group size. Therefore, the actual size differed between groups (Supplementary Figure 3, Supplementary table 1). Data collection and analysis were not performed blind to the conditions of the experiments.

### 3.3 The multi-player three-armed bandit task

To study the relationship between social information use and collective behavioural dynamics, we focused on a well-established learning-and-decision problem called a ‘multi-armed bandit’ task, represented here as repeated choices between three slot machines (Supplementary Figure 1, Video 1, for detail see Supplementary Method). Participants played the task for 70 rounds. The slots paid off money noisily (in the US cents), varying around two different means during the first 40 rounds such that there was one ‘good’ slot and two other options giving poorer average returns. From the round 41st, however, one of the ‘poor’ slots abruptly increased its mean payoff to become ‘excellent’ (i.e. superior to ‘good’). The purpose of this environmental change was to observe the effects of maladaptive herding by potentially trapping groups in the out-of-date suboptimal (good) slot, as individuals did not know whether or how an environmental change would occur. Through making choices and earning a reward from each choice, individuals could gradually learn which slot generated the highest rewards.

In addition to this asocial learning, we provided social information for each member of the group specifying the frequency with which group members chose each slot. All group members played the same task with the same conditions simultaneously, and all individuals had been instructed that this was the case, and hence understood that the social information would be informative.

Task uncertainty was experimentally manipulated by changing the difference between the mean payoffs for the slot machines. In the task with the least uncertainty, the distribution of payoffs barely overlapped, whilst in the task with the greatest uncertainty the distribution of payoffs overlapped considerably (Supplementary Figure 1).

### 3.4 The computational learning-and-decision-making model

We modelled individual behavioural processes by assuming that individual *i* makes a choice for option *m* at round *t*, in accordance with the choice-probability *P_i,t_*(*m*) that is a weighted average of social and asocial influences:

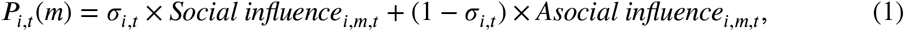

where *σ_i_*_,*t*_is the *social learning weight* (0 ≤ *σ_i_*_,*t*_ ≤ 1).

For the social influence, we assumed a frequency-dependent copying strategy by which an individual copies others’ behaviour in accordance with the distribution of social frequency information ^49–51;55^:

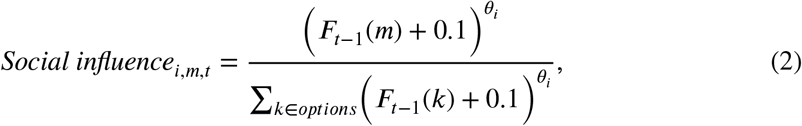

where *F_t_*_−1_(*m*) is a number of choices made by other individuals (excluding her/his own choice) for the option *m* in the preceding round *t* − 1 (*t* ≥ 2). *0* is individual *i*’s *conformity exponent*, −∞ ≤ *θ_i_* ≤ +∞. When this exponent is larger than zero, higher social influence is given to an option which was chosen by more individuals (i.e. positive frequency bias). When this exponent is larger than zero (*θ_i_ >* 0), higher social influence is afforded to an option chosen by more individuals (i.e. positive frequency bias), with conformity bias arising when *θ_i_ >* 1, such that disproportionally more social influence is given to the most common option ^28^. When *θ_i_ <* 0, on the other hand, higher social influence is afforded to the option that fewest individuals chose in the preceding round *t* − 1 (i.e. negative frequency bias). To implement the negative frequency dependence, we added a small number 0.1 to *F* so that an option chosen by no one (i.e. *F_t_*_−1_ = 0) could provide the highest social influence when *θ_i_ <* 0. Note, there is no social influence when *θ_i_* = 0 because in this case the ‘social influence’ favours an uniformly random choice, i.e., 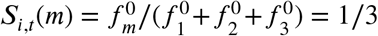, independent of the social frequency distribution.

Note also that, in the first round *t* = 1, we assumed that the choice is only determined by the asocial softmax function because there is no social information available.

For the asocial influence, we used a standard reinforcement learning with ‘softmax’ choice rule ^75^, widely applied in human social learning studies e.g. ^35;51;55^. An individual *i* updates the estimated average reward associated with an option *m* at round *t*, namely Q-value (*Q_i,t_*(*m*)), according to the Rescorla-Wagner rule as follows:

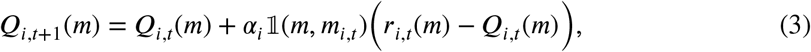

where *α_i_* (0 ≤ *α_i_* ≤ 1) is a *learning rate* parameter of individual *i* determining the weight given to new experience and *r_i_*_,*t*_(*m*) is the amount of monetary reward obtained from choosing the option *m* in round 𝟙. (*m*, *m_i,t_*) is the binary action-indicator function of individual *i*, given by

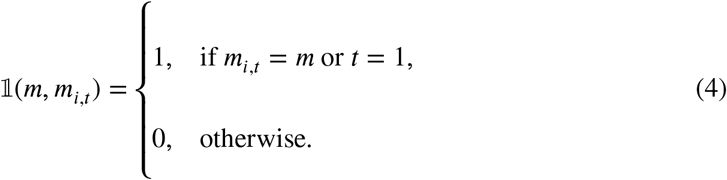

Therefore, *Q_i_*_,*t*_(*m*) is updated only when lIthe option *m* was chosen; when the option *m* was not chosen, *Q_i_*_,*t*_(*m*) is not updated (i.e. *Q_i_*_,*t*+1_(*m*) = *Q_i_*_,*t*_(*m*)). Note that, only in the first round *t* = 1, all Q-values are updated by using the chosen option’s reward *r_i_*_,1_(*m*), so that the individual can set a naive ‘intuition’ about the magnitude of reward values she/he would expect to earn from a choice in the task; namely, *Q_i_*_,*t*=2_(1) = *Q_i_*_,*t*=2_(2) = *Q_i_*_,*t*=2_(3) = *α_i_r_i_*_,*t*=1_(*m*). In practical terms, this prevents the model from being overly sensitive to the first experience. Before the first choice, individuals had no prior preference for either option (i.e. *Q_i_*_,1_(1) = *Q_i_*_,1_(2) = *Q_i_*_,1_(3) = 0).

The Q-value is then translated into the asocial influence through the softmax (or logit choice) function:

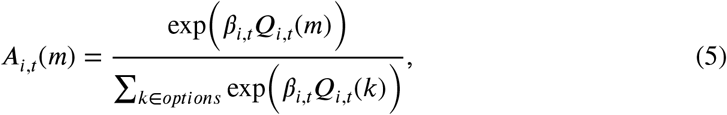

where *β_i_*_,*t*_, called *inverse temperature*, manipulates individual *i*’s sensitivity to the Q-values (in other words, controlling the proneness to explore). As *β_i_*_,*t*_ goes to zero, asocial influence approximates to a random choice (i.e. highly explorative). Conversely, if *β_i_*,_*t*_→ +∞, the asocial influence leads to a deterministic choice in favour of the option with the highest Q-value (i.e. highly exploitative). For intermediate values of *β_i_*,_*t*_, individual *i* exhibits a balance between exploration and exploitation ^35;68^. We allowed for the possibility that the balance between exploration-exploitation could change as the task proceeds. To depict such time dependence in exploration, we used the equation: 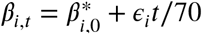. If the slope *ϵ_i_* is positive (negative), asocial influence *A_i_*,_*t*_becomes more and more exploitative (explorative) as round *t* increases. For a model fitting purpose, the time-dependent term *ϵ_i_t* is scaled by the total round number 70.

We allowed that the social learning weight *σ_i_*_,*t*_could also change over time as assumed in the inverse temperature *β_i_*,*_t_*. To let *σ_i_*,_*t*_ satisfy the constraint 0 ≤ *σ_i_*,_*t*_ ≤ 1, we used the following sigmoidal function:

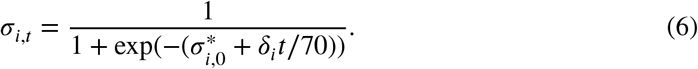

If the slope *δ_i_* is positive (negative), the social influence increases (decreases) over time. We set the social learning weight equal to zero when group size is one (i.e. when an individual participated in the task alone and/or when 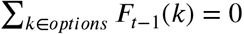.

We modelled both the inverse temperature *β_i_*,_*t*_and the social learning weight *σ_i_*_,*t*_as a time function since otherwise it would be challenging to distinguish different patterns of learning in this social learning task ^63^. The parameter recovery test confirmed that we were able to differentiate such processes under these assumptions (Supplementary Figure 7, Supplementary Figure 8). While we also considered the possibility of the conformity exponent being time-dependent (i.e. 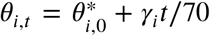, the parameter recovery test suggested that the individual slope parameter *y_i_* was not reliably recovered (Supplementary Figure 9), and hence we concentrated our analysis on the time-independent *θ_i_* model. We confirmed that instead using the alternative model where both social learning parameters were time-dependent (i.e. *σ_i_*_,*t*_and *θ_i_*_,*t*_) did not qualitatively change our results (Supplementary Figure 10).

One concern might be the asymmetry between the asocial softmax influence which takes many prior experiences into account (depending upon a learning rate) and the social influence referring only the most recent frequency information *F_t_*_−1_. The choice frequency appeared at round *t* is the most reliable social information, compared to the past frequencies, because it could be the most ‘updated’ information as long as the other individuals have made informed decisions in their best knowledge. In contrast, option’s reward just obtained at *t* − 1, which was independently and randomly drawn from a probability distribution, is less reliable than accumulated Q-values taking past rewards into account. Although many other formulations for asocial and social learning processes were possible, we believe that our current choice – time-depth asocial reinforcement learning with the most-updated-frequency-dependent copying was a reasonable first step.

In summary, the model has six free parameters that were estimated for each individual human participant; namely, *a_i_*, 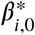, *ϵ_i_*, 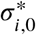, *δ_i_*, and *θ_i_*. To fit the model, we used a hierarchical Bayesian method (HBM), estimating the global means (*μ_a_*, 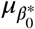, *μ_ϵ_*, 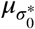, *μ_δ_*, and *μ_θ_*) and the global variations (ν_*a*_, 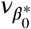, *v_ϵ_*, 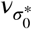, ν_*δ*_, and ν_*θ*_) for each of the three experimental conditions (i.e. the Low-, Moderate-and High-uncertain condition), which govern overall distributions of individual parameter values. It has become recognised that the HBM can provide more robust and reliable parameter estimation than conventional maximum likelihood point estimation in complex cognitive models ^76^, a conclusion with which our parameter recovery test agreed (Supplementary Figure 7, Supplementary Figure 8).

### 3.5 Agent-based model simulation

We ran a series of individual-based model simulations assuming that a group of individuals play our three-armed bandit task for 90 rounds (under the Moderate-uncertainty condition) and that individuals behave in accordance with the computational learning-and-decision model. We varied the group size (*n* ∈ {3, 10, 30}), the mean social learning weight (*σ̄* ∈ {0.01, 0.1, 0.2, 0.3, …, 0.9}) and the mean conformity exponent (*θ̄* ∈ {0.5, 1, 3, 6}), running 10,000 replications for each of the possible parameter × group size combinations. As for the other parameter values (e.g. the asocial reinforcement learning parameters; *α*, 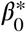, *ϵ*), here we used the experimentally fitted global means (Table 2 and Supplementary table 3). Relaxation of this assumption (i.e. using a different set of asocial learning parameters) does not qualitatively change our story (Supplementary Figure 2). Note that each individual’s parameter values were randomly drawn from the distributions centred by the global mean parameter values fixed to each simulation run. Therefore, the actual composition of individual parameter values were different between individuals even within the same social group.

### 3.6 Generalised linear mixed models

To directly analyse the effects of group size and task uncertainty on the time evolution of decision performance, we conducted a statistical analysis using a phenomenological model, namely, a hidden Markov process logistic regression without assuming any specific learning-and-decision-making processes. The dependent valuable was whether the participant chose the best option (1) or not (0). The model includes fixed effects of grouping *ξ*, standardised group size *ω*, and an intercept with a random effect of individuals *μ* + *p_i_*. We assumed that the intercept and the effect of grouping change from round to round, as a random walk process. For the effect of group size we considered the effect of the 1st environment 1 ≤ *t* ≤ 40 and that of the 2nd environment, namely, *ω*_1_ and *ω*_2_, separately.

To examine whether increasing group size and increasing task uncertainty affected individual use of the positive frequency-dependent copying strategy, we used a hierarchical Bayesian logistic regression model with a random effect of groups. The dependent valuable was whether the participant used the positive frequency-dependent copying (1) or not (0). The model includes fixed effects of group size (standardised), task uncertainty (0: Low, 0.5: Moderate, 1: High), age (standardised), gender (0: male, 1: female, NA: others or unspecified), and possible two-way interactions between these fixed effects.

We also investigated the effects of both group size and the task’s uncertainty on the fitted values of the learning parameters. We used a hierarchical Bayesian gaussian regression model predicting the individual fitted parameter values. The model includes effects of group size (standardised), task uncertainty (0: Low, 0.5: Moderate, 1: High), age (standardised), gender (0: male, 1: female, NA: others or unspecified), and two-way interactions between these fixed effects. We assumed that the variance of the individual parameter values might be contingent upon task uncertainty because we had found in the computational model-fitting result that the fitted global variance parameters (i.e. 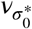, ν_*δ*_, and ν_*θ*_) were larger in more uncertain conditions (Supplementary table 2).

### 3.7 Statistical analysis

We used a hierarchical Bayesian method (HBM) to estimate the free parameters of our statistical models, including both the phenomenological regression model and the computational learning-and-decision-making model. The HBM allows us to estimate individual differences, while ensures these individual variations are bounded by the group-level global parameters. The HBM was performed under Stan 2.16.2 (http://mc-stan.org) in R3.4.1 (https://www.r-project.org) software. The models contained at least 4 parallel chains and we confirmed convergence of the MCMC using both the Gelman-Rubin statistics and the effective sample sizes. Full details of the model fitting procedure and prior assumptions are shown in the appendix.

#### 3.7.1 Parameter recovery test

To check the validity of our model-fitting method, we conducted a ‘parameter recovery test’ so as to examine how well our model fitting procedure had been able to reveal true individual parameter values. To do this, we generated synthetic data by running a simulation with the empirically fitted global parameter values, and then re-fitted the model with this synthetic data using the same procedure. The parameter recovery test showed that the all true global parameter values were fallen into the 95% Bayesian credible interval (Supplementary Figure 7), and at least 93% of the true individual parameter values were correctly recovered (i.e. 96% of *α_i_*, 93% of 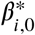, 95% of *ϵ_i_*, 97% of 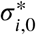, 96% of *δ_i_* and 97% of *θ_i_* values were fallen into the 95% Bayesian CI. Supplementary Figure 7).

#### 3.7.2 Categorisation of individual learning strategies

Based on the 50% CI of the individual *conformity exponent* parameter values *0_i_*, we divided the participants into the following three different social learning strategies. If her/his 50% CI of *θ_i_* fell above zero (*θ_lower_* > 0), below zero (*θ_upper_* < 0) or including zero (*θ_lower_* ≤ 0 ≤ *θ_upper_*), she/he was categorised as a ‘positive frequency-dependent copier’, a ‘negative frequency-dependent copier’, or a ‘random choice individual’, respectively. We used the 50% Bayesian CI to conduct this categorisation instead of using the more conservative 95% CI because the latter would cause much higher rates of ‘false negatives’, by which an individual who applied either a positive frequency-dependent copying or a negative-frequency dependent copying strategy was falsely labelled as an asocial random choice individual (Supplementary Figure 7). Four hundred agents out of 572 (≈ 70%) were falsely categorised as a random choice learner in the recovery test when we used the 95% criterion (Supplementary Figure 7). On the other hand, the 50% CI criterion seemed to be much better in terms of the false negative rate which was only 18.5% (i.e. 106 agents), although it might be slightly worse in terms of ‘false positives’: Thirty-seven agents (6.5%) were falsely labelled as either a positive frequency-dependent copier or a negative-frequency dependent copier by the 50% CI, whereas the false positive rate of the 95% CI was only 0.2% (Supplementary Figure 7). To balance the risk of false positives and false negatives, we decided to use the 50% CI which seemed to have more strategy detecting power.

#### 3.7.3 The post-hoc model simulation

So as to evaluate how accurately our model can generate observed decision pattern in our task setting, we ran a series of individual-based model simulation using the fitted individual parameter values (i.e. means of the individual posterior distributions) for each group size for each uncertainty condition. At the first step of the simulation, we assigned a set of fitted parameters of a randomly-chosen experimental subject from the same group size and the same uncertain condition to an simulated agent, until the number of agents reaches the simulated group size. We allowed duplicate choice of experimental subject in this parameter assignment. At the second step, we let this synthetic group of agents play the bandit task for 90 rounds. We repeated these steps 10,000 times for each group size, task uncertainty.

### 3.8 Data availability

Both experimental and simulation data are available on an online repository (https://github.com/WataruToyokawa/ToyokawaWhalenLaland2018).

### 3.9 Code availability

The browser based online task was built by Node.js (https://nodejs.org/en/) and socket.io (https://socket.io), and the code are available on a GitHub repository (https://github.com/WataruToyokawa/MultiPlayerThreeArmedBanditGame). Analyses were conducted in R (https://www.r-project.org) and simulations of individual based models were conducted in Mathematica (https://www.wolfram.com), both are available on an online repository (https://github.com/WataruToyokawa/ToyokawaWhalenLaland2018).

## 5 Authors’ contributions

WT, AW and KNL planned the study and built the computational model. WT ran simulations. WT and AW made the experimental material, ran the web-base experiment, and collected the experimental data. WT, AW and KNL analysed the data and wrote the manuscript.

## 6 Acknowledgements

The experiment was supported by The John Templeton Foundation (KNL; 40128) and Suntory Foundation Research Support (WT; 2015-311). The computer simulations and computational model analyses were supported by JSPS Overseas Research Fellowships (WT; H27-11). The phenomenological model analyses were supported by JSPS KAKENHI Grant Number 17J01559. The funders had no role in study design, data collection and analysis, decision to publish or preparation of the manuscript.

