## Supplementary Materials for "Social learning strategies regulate the wisdom and madness of interactive crowds"

### 1 Supplementary Methods

#### 1.1 Amazon Mechanical Turk

The online task was advertised as a 'HIT (Human Intelligence Task)' entitled 'Lottery Choice Experiment (about 15 mins + fun + bonus!)' in Amazon Mechanical Turk (AMT). The HIT was only available for individuals whose 'HIT Approval Rate' was greater than or equal to 90% and who live in the US.

On the advertisement page, we stressed that there could be extra payoff to subjects depending on their performance in the task; that stability of their Internet connection was necessary; and that they could participate in the task only once (Supplementary Video 1).

At the bottom of the advertisement page, an URL link to our experimental server would appear. The link was not shown until the participants decided to join the task. There were also text forms into which the participants could input their own confirmation code, which they would get by finishing the experimental task, and where they could add comments on the task. By clicking the submit button below they could complete the task, allowing the monetary reward to be paid through Amazon system.

#### 1.2 A written consent form and an instruction for the task

On clicking the URL link shown at the bottom of the HIT advertisement page, participants proceeded to a consent form that emphasised data anonymity and asked them not to interact with anyone during the task (Supplementary Video 1). Scrolling this consent form page, participants proceeded to sign the form. After answering all the YES/NO questions and inputting the

CAPTCHA, participants could proceed to instructions.

On the first page of the instructions participants were informed that the study was split into two parts: an interactive economic decision-making game and a short survey. On the second page of the instructions the details of the decision task were explained with illustrations.

Full details and code of the task are available in GitHub (https://github.com/WataruToyokawa/MultiPlayerThreeArmedBanditGame).

#### 1.3 Participants in the online experiment

A total of 755 subjects (354 females, 377 males, 2 others and 22 unspecified; mean age (1 $s.d.$) = 34.33 (10.9)) participated in our incetivised economic behavioural experiment (Figure S2). The experimental sessions were conducted in December 2015 and in January 2016. We excluded subjects who disconnected to the online task before completing at least the first 30 rounds from our learning model fitting analysis, resulted in 699 subjects (573 subjects entered the group (i.e. $n \geq 2$) condition and 126 entered the solitary (i.e. $n = 1$) condition). The task was advertised using Amazon's Mechanical Turk (AMT; https://www.mturk.com; see Supplementary Video 1; Supplementary Video 3), so that the participants could enter anonymously through their own internet browser window. Upon connecting to the experimental game web page, the participants might be required to wait on other participants at the virtual 'waiting room' for up to 5 minutes or until the requisite number of participants arrived, whichever was sooner, before the task starts. The participants were payed 25 cents for a show-up fee plus a waiting-bonus at a rate of 12 cents per minute (i.e. *pro rata* to 7.2 USD per hour) and a game bonus ($mean \pm 1 s.d. = 1.7 \pm 0.79$ USD) depending on their performance in the task. The total time, including net time spent in the waiting room, tended to be less than 10 minutes.

#### 1.4 The multi-player three-armed bandit task

The participants performed a three-armed bandit task for 70 rounds. Each round started with the choice stage at which three slot machines appeared on the screen (Supplementary Figure 1; Supplementary Video 2). Participants chose a slot by clicking the mouse pointer (or tapping it if they used a tablet computer). Participants had a maximum of 8 seconds to make their choices. If no choice was made during the choice stage, a 'TIME OUT' message appeared in the centre of the screen without a monetary reward (average number of missed rounds per participant was 0.18 out of 70 rounds). Participants were able to know the rest of the choice time by seeing a 'count-down bar' shown at the top of the experimental screen.

Each option yielded monetary rewards randomly drawn from a normal probability distribution unique to each slot, rounded up to the next integer, or truncated to zero if it would have been a negative value (Supplementary Figure 1). The standard deviations of the probabilistic payoff distributions were identical for all slots and did not change during the task (the $s.d. = 0.55$; although it actually was slightly smaller than 0.55 due to the zero-truncation). The mean values of the probabilistic pay-off were different between the options. 'Poor', 'good' and 'excellent' slots generated the lowest, intermediate and the highest rewards on average, respectively. In the first 40 rounds, there were two poor and one good options. After the round 40th, one of the poor option abruptly changed to an excellent option (i.e. environmental change), and from the 41st round there were poor, good and excellent options. The purpose of this environmental change was to observe the effects of inflexible herding by potentially trapping groups in the out-of-date suboptimal (good) slot, as individuals did not know whether or how an environmental change would occur. Through making choices and earning a reward from each choice, individuals could gradually learn which slot generated the highest rewards.

Once all the participants in the group made a choice (or had been time-outed), they proceeded to the feedback stage in which they could see their own payoff from the current choice for two seconds ('0' was shown if they had been time-outed), while they could not see others' reward values. After this feedback stage, subjects proceeded to the next round's choice stage. From the second round, a distribution of choices made by all participants in the group at the preceding round (i.e. the social frequency information) was shown below each slot.

Before the task started, participants had read an illustrated instruction which told them that they would play 70 rounds of the task, that the payoff would be randomly generated per choice but associated with a probability distribution unique to each slot machine, i.e. the profitability of the slot might be different from each other, that the environment might change during the task so that the mean payoff from the slots might secretly change during the task, and that their total payout were decided based on the sum of all earnings they achieved in the task. We also explicitly informed subjects that all participants in the same group played the identical task so that they could infer that the social information was informative. However, we did not specify either the true mean payoff values associated with each option, or when and how the mean payoff would actually change. After reading these instructions, participants proceeded to a 'tutorial task' without any monetary reward and without the social frequency information, so as to become familiar with the task.

After they completed the behavioural task or were excluded from the task due to a bad internet connection or due to opening another browser window during the task (see the 'Reducing the risk of cheating' section in the supplementary material), subjects proceeded to a brief questionnaire page asking about demographic information, which were skippable. Finally, the result screen was shown, informing the total monetary reward she/he earned as well as a confirmation code unique for each participant. Participants could get monetary reward through Amazon Mechanical Turk (AMT) by inputting the confirmation code into the form at the AMT's task page (see Supplementary material for further details of experimental procedure).

##### 1.5 Reducing the risk of cheating

To minimise the risk of multiple accesses from the same person, we introduced the restriction that a single 'worker ID' associated with participants' AMT accounts, could participate only once in the experiment. We rejected access from the same IP address: If a participant's IP address had already been stored in our database, the participant directly proceeded from the instruction page to the questionnaire page. In that case, 25 cents show-up fee was still paid because it was possible that different persons might use the same IP address.

To minimise the risk of opening other browser windows during the task (for example, browsing other websites), we used 'Page Visibility API' (https://developer.mozilla.org/en-US/docs/Web/API/Page_Visibility_API) to track whether the experimental browser window was always active and not hidden by other browser windows or tabs for more than 1 second. If it was detected that the experimental window was in a hidden state, the participant was automatically redirected to the questionnaire page. In that case, 25 cents show-up fee plus a waiting-bonus (if applicable) and a game-bonus earned so far were paid. In the instruction, participants were warned not to open any other browser windows/tabs during the task and were informed that they would be eliminated from the session if they did so.

#### 1.6 Hierarchical Bayesian parameter estimation

We used hierarchical Bayesian method (HBM) to estimate the free parameters of our learning model. HBM allows us to estimate individual differences, while this individual variation is bounded by the group-level (i.e. hyper) parameters. To perform HBM, we used Stan 2.16.2 in R 3.4.1.

In our model, there are 6 individual parameters; namely, $\alpha_i$, $\beta^*_{i,0}$, $\epsilon_i$, $\sigma^*_{i,0}$, $\delta_i$, and $\theta_i$. Because the learning rate $\alpha_i$ is bounded between 0 and 1, we estimated $\alpha^*_i$ rather than $\alpha_i$ itself ($-\infty \leq \alpha^*_i \leq +\infty$), which is given by the following sigmoidal function:

$$\alpha_i = \frac{1}{1 + \exp(-\alpha^*_i)}. \tag{1}$$

We assumed the Student's t distributions for individual random effects of each parameter so as to allow a few 'outliers', because the Student's t distribution has a longer tail compared to a normal distribution. To do so, we used the following reparameterization for each $parameter(.) \in \{\alpha^*_i, \beta^*_{i,0}, \epsilon_i, \sigma^*_{i,0}, \delta_i, \theta_i\}$:

$$parameter(.)_i = \mu_{(.),c} + \nu_{(.),c} * parameter(.)\_raw_i, \tag{2}$$

where $\mu_{(.),c}$ is a global mean of the $parameter(.)$ in the condition $c$ ($c \in$ {Low- , Moderate-, High-uncertainty condition}), and $\nu_{(.),c}$ is a global scale parameter of the individual variations in condition $c$, which is multiplied by a standardised individual random variable $parameter(.)\_raw_i$ drawn from

$$parameter(.)\_raw_i \sim Student\_t\big(df = 4, location = 0, scale = 1\big). \tag{3}$$

As for the global parameters, we used a normal and a half-normal prior distributions for $\mu_{(.),c}$ and $\nu_{(.),c}$, respectively:

$$\mu_{(.),c} \sim Normal\big(mean = 0, sd = 5\big), \tag{4a}$$

$$\nu_{(.),c} \sim Normal^{+}\big(mean = 0, sd = 3\big). \tag{4b}$$

In summary, there are 36 global free parameters (= 6 $\mu$s and 6 $\nu$s for 3 different conditions each). A total of 2000 iterations were performed after 1000 warm-up with thin = 5 for each of 8 chains (= 2000 samples / 5 steps × 8 chains = a total of 3200 samples). We used the Gelman-Rubin statistics (as known as $\hat{R}$) as well as the effective sample sizes (ESS) so as to check the convergence of the MCMC samples. All global parameter values had $\hat{R} \approx 1.00 \leq 1.10$ indicating that chains are converged to the target distributions. The ESS of model parameters were typically greater than 500 (out of 3200 total samples).The minimum ESS of global-parameters was 233 (on $\nu_{\epsilon,Low}$). Visual inspection of the parameters with smaller ESSs confirmed their convergence to target distributions. We confirmed that changing both *df* (i.e. broadness of the tail) of the Student's t prior distributions and *sd* of the Normal prior distributions did not change our findings.

#### 1.7 Parameter recovery test

To assess the adequacy of the hierarchical Bayesian model-fitting method, we tested how well the HBM could recover 'true' parameter values that were used to simulate synthetic data. We simulated participants' behaviour assuming that they behave according to the model with each parameter setting. We generated 'true' parameter values for each simulated agent based on the experimentally fit global parameters (Table 1 in the main text). We then simulated synthetic behavioural data and recovered their parameter values using the HBM described above.

#### 1.8 Time-dependent conformity exponent $\theta_{i,t}$ model

We also considered the possibility of the conformity exponent being time-dependent (i.e. $\theta_{i,t} = \theta^*_{i,0} + \gamma_i t/70$). If the slope $\gamma_i$ is positive (negative), the frequency-dependent bias increases (decreases) over time. In this model, there are 7 individual parameters; namely, $\alpha_i$, $\beta^*_{i,0}$, $\epsilon_i$, $\sigma^*_{i,0}$, $\delta_i$, $\theta^*_{i,0}$ and $\gamma_i$. We fitted this model to the experimental data using the HBM descried above.

#### 1.9 A hidden Markov model regression

To analyse effects of group size and task uncertainty on the decision performance, we conducted a statistical analysis using a phenomenological model, namely, a hidden Markov process logit model (eq. 5). The dependent valuable was whether the participant chose the best option (1) or not (0). The individual $i$'s probability of choosing the best option at time $t$ under the uncertainty condition $c$, namely $Prob_{c,i,t}$, depends basically on a global mean $\mu_{c,t}$ and the random effect of individuals $\rho_i$. If the individual $i$ plays with other individual(s) (i.e. if the binary indicator of group condition $\mathbb{1}_g = 1$), the probability $Prob_{c,i,t}$ is also contingent upon an effect of grouping

$\left\{\xi_{c,t} + \omega_{1\text{ or }2} \times \text{standardised group size}\right\}$. We considered the group size effects $\omega$ for the 1st and 2nd environments separately; namely, $\omega_1$ and $\omega_2$.

$$
\begin{aligned}
\text{Choosing the best option or not} &\sim Bernoulli\left(Prob_{c,t}\right), \qquad (5)\\
\text{logit}\left(Prob_{c,i,t}\right) &= \begin{cases} \mu_{c,t} + \rho_i + \mathbb{1}_g \left\{\xi_{c,t} + \omega_1 \times \text{group size}\right\}, & \text{if } 1 \leq t \leq 40, \\ \mu_{c,t} + \rho_i + \mathbb{1}_g \left\{\xi_{c,t} + \omega_2 \times \text{group size}\right\}, & \text{if } t \geq 41, \end{cases}\\
\mu_{c,t} &= \mu_{c,t-1} + \text{Cauchy}\left(0, \sigma_\mu\right),\\
\xi_{c,t} &= \xi_{c,t-1} + \text{Cauchy}\left(0, \sigma_\xi\right).
\end{aligned}
$$

Note that both $\mu$ and $\xi$ are determined by a Markov process, moving randomly under a Cauchy probability distribution. The Cauchy distribution has a longer tail than a Gaussian distribution, allowing a sudden 'jump' in the random walk which should occur at $t = 41$ in our task.

We have also modelled participants' actual monetary reward as a performance measurement, using the same model as eq. 5 except the following parts:

$$
\begin{aligned}
\text{Payoff earned} &\sim Normal^{+}\left(Mean_{c,t}, \sigma_m\right), \qquad (6)\\
Mean_{c,i,t} &= \begin{cases} \mu_{c,t} + \rho_i + \mathbb{1}_g \left\{\xi_{c,t} + \omega_1 \times \text{group size}\right\}, & \text{if } 1 \leq t \leq 40, \\ \mu_{c,t} + \rho_i + \mathbb{1}_g \left\{\xi_{c,t} + \omega_2 \times \text{group size}\right\}, & \text{if } t \geq 41. \end{cases}
\end{aligned}
$$

We estimated free parameters: $\mu_{c,t}$, $\xi_{c,t}$, $\omega_1$ and $\omega_2$, using the same MCMC procedure we used for our computational mechanism model (Section 2 in this supplementary method).

### 2 Supplementary Notes

#### 2.1 Individual-based simulation using other parameter sets

Individual-based model simulations using a different set of asocial learning parameters suggest that our main findings from the simulation (Figure 1, 2 in the main text) are broadly robust in a range of parameter combinations (Supplementary Figure 2).

#### 2.2 Results of a time-dependent conformity model ($\theta_{i,t}$)

##### 2.2.1 Parameter recovery test

The parameter recovery test showed that the all true global parameter values were fallen into the 95% Bayesian credible interval (Supplementary Figure 9). Correlations between individual true parameters and recovered parameters were all positive, while the correlation coefficients of both $\theta^*_{i,0}$ and $\gamma_i$ were lower than other parameters (Supplementary Figure 9). At least 89% of the true individual parameter values were correctly recovered (i.e. 97% of $\alpha_i$, 96% of $\beta^*_{i,0}$, 97% of $\epsilon_i$, 96% of $\sigma^*_{i,0}$, 94% of $\delta_i$, 96% of $\theta^*_{i,0}$ and 89% of $\gamma_i$ were fallen into the 95% Bayesian CI).

##### 2.2.2 Fitting to our experimental data

In order to compared the findings from the time-independent $\theta_i$ model (i.e. the model used in the main text), we again categorized the participants as deploying three different learning strategies based on their mean fitted conformity exponent values; namely, the 'positive frequency-dependent copying' strategy ($\bar{\theta}_i \gg 0$), the 'negative-frequency dependent copying' strategy ($\bar{\theta}_i \ll 0$) and the 'random choice' strategy ($\bar{\theta}_i \approx 0$). Note, the conformity exponent here is averaged over time: $\bar{\theta}_i = (\sum_t \theta_{i,t})/70$. Supplementary Figure 10 suggests that the patterns were consistent with Figure 4 in the main text, and hence our conclusion was not changed.

Individual frequency dependence changed over time (Supplementary Figure 10B). The conformity exponents generally increased with experimental round, while some individuals in the High-uncertain conditions decreased rather than accelerated their frequency dependence over time. However, note that the fitting of slope parameter $\gamma_i$ was relatively unreliable (i.e. only 89% of individual parameters were recovered correctly). Extensive variation in both the social learning weigh $\sigma_{i,t}$ and the conformity exponent $\theta_{i,t}$ found in high-uncertain circumstances are consistent with the main findings (Figure 5).

### 3 Supplementary Figures

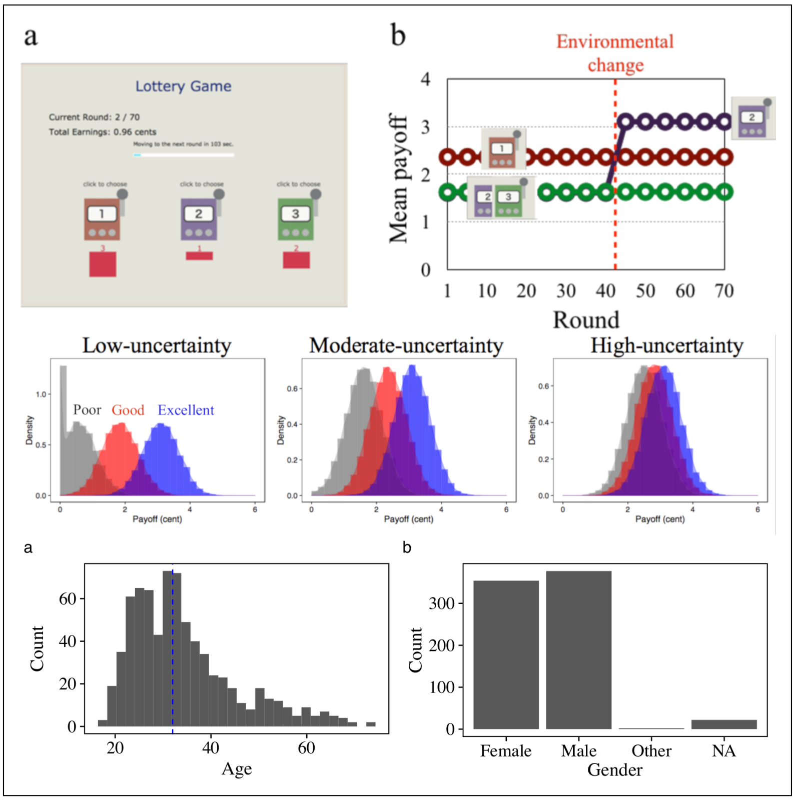


**Figure 1. The web-based three-armed bandit experimental task**. (Upper row, a) Participants could choose an option by clicking one of the slot machines. The frequency distribution of choices made by participants in the same group in the preceding round (i.e. the social frequency information) is shown by red numbers below each option. (Upper row, b) Example of mean payoffs for each option in the task. The association between each option's number and its payoff was randomly assigned across experimental sessions. (Middle row) The distributions of payoffs generated by each of the slot machines for each condition. The poor, good and excellent slot are indicated by grey, red and blue, respectively. The payoff was truncated to zero if it would have been a negative value. (Lower row) Histograms of the participants' age and gender. Note that these data were inputted by participants themselves on the questionnaire forms.

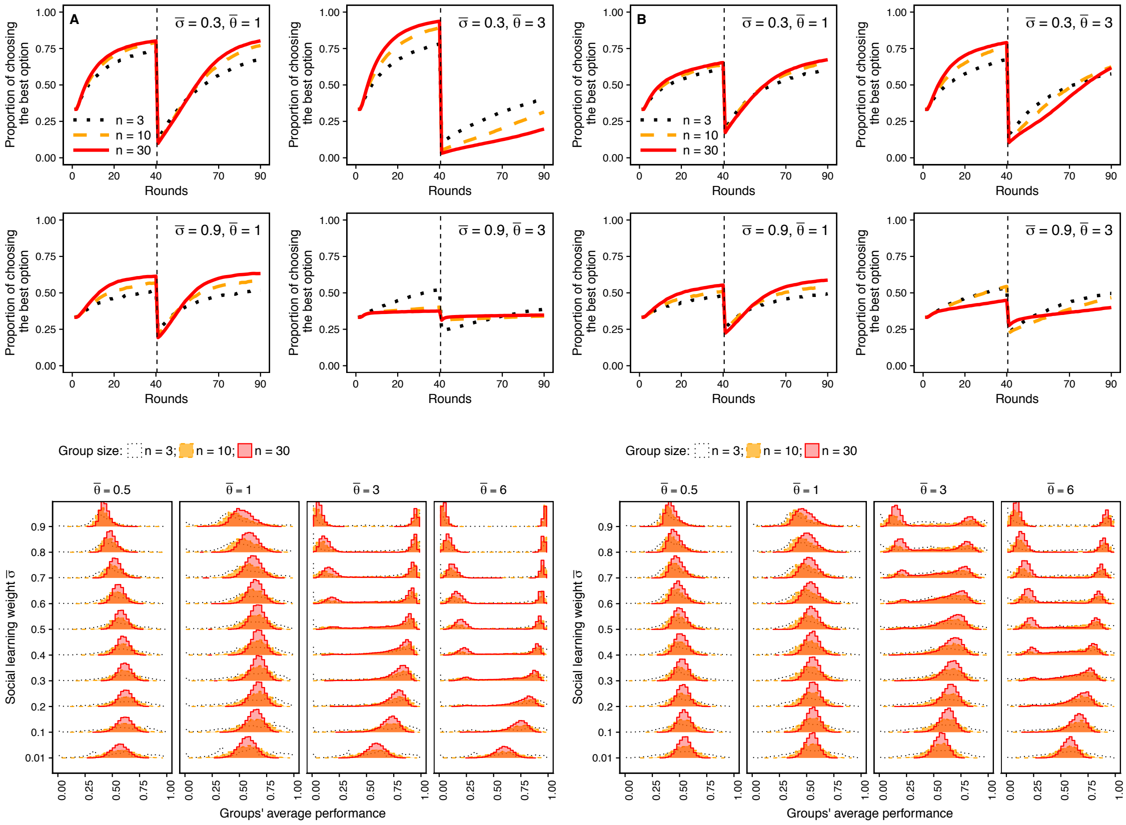

**Figure 2. The same figure as Fig. 1 and Fig. 2 in the main text, except for the asocial learning parameter settings**. (Left column A): $\mu_\alpha = 0.7$, $\mu_{\beta_0^*} = 2$, $\mu_\epsilon = 4$, $\nu_\alpha = 1$, $\nu_{\beta_0^*} = 1$, $\nu_\epsilon = 1$, $\nu_\sigma = 1$, and $\nu_\theta = 1$. (Left column B): $\mu_\alpha = 0.8$, $\mu_{\beta_0^*} = 0.5$, $\mu_\epsilon = 3$, $\nu_\alpha = 1$, $\nu_{\beta_0^*} = 2$, $\nu_\epsilon = 2$, $\nu_\sigma = 2$, and $\nu_\theta = 2$

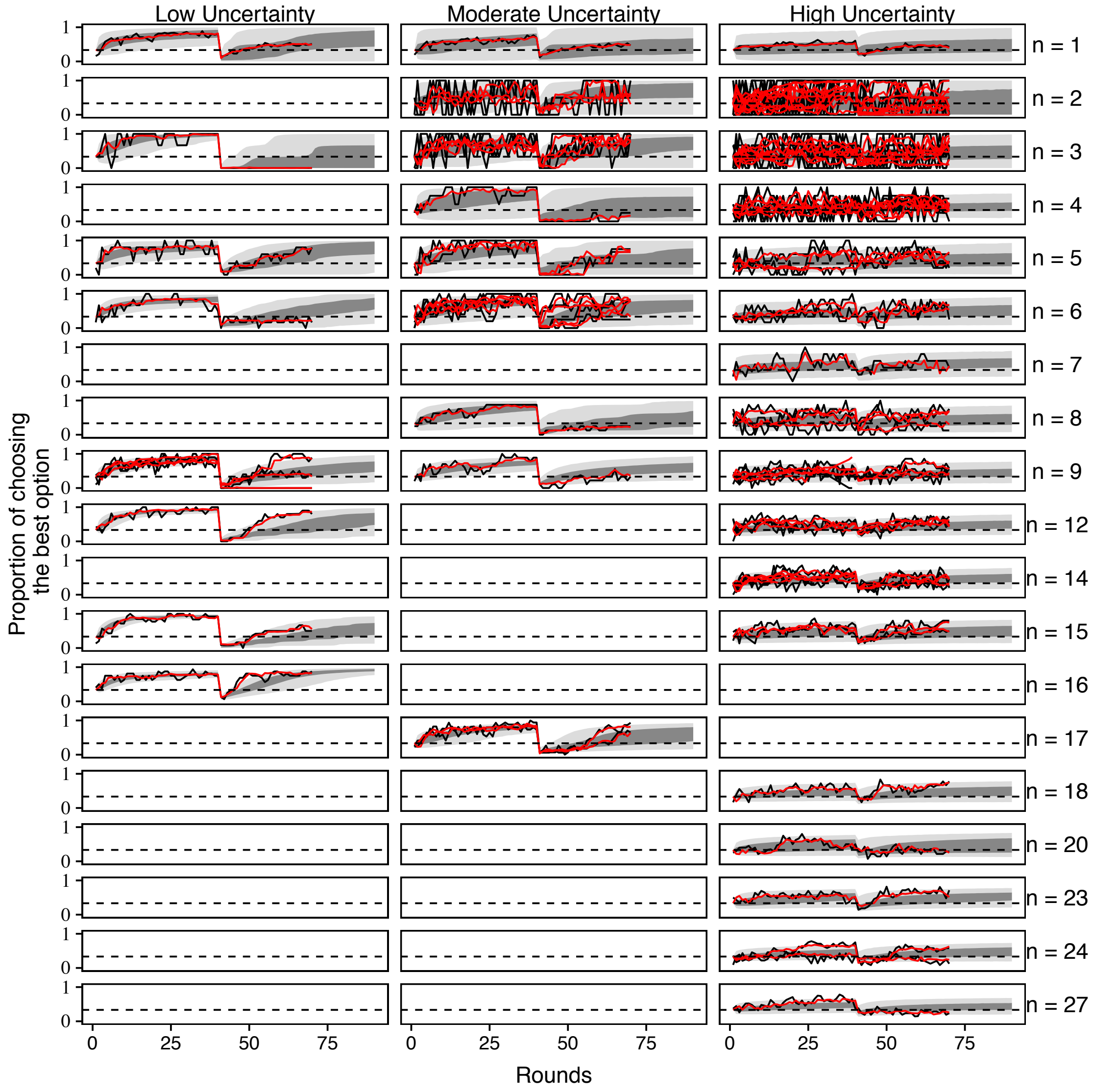


**Figure 3. Dynamics of each group's mean performance (black lines) and fitted dynamics derived from the learning-and-decision-making model (red lines)**. Group size is indicated in the righthand side. The 95% and 50% quantiles of trajectories of the post-hoc simulation are shown as light and dark shades, respectively. The horizontal dashed line indicates a chance-level (i.e. $1/3$). Note that the experimental task finished at the 70th round, whereas the simulation ran for 90 rounds.

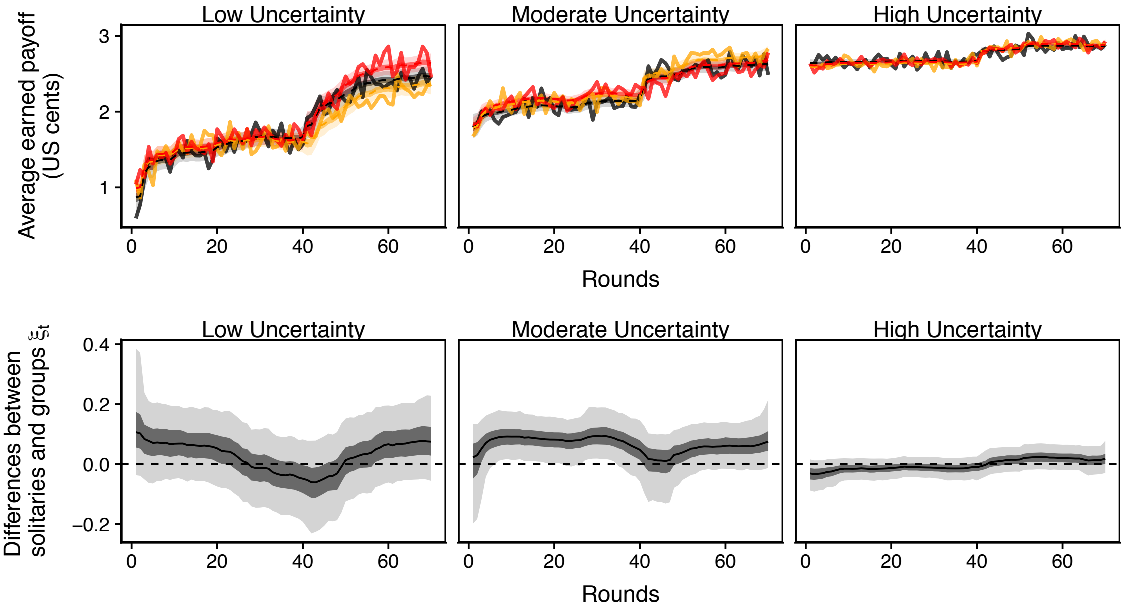


**Figure 4. A Gaussian model regression for the participants' payoff**. (Top row): The average earnings (in US cents) of the experimental participants (red: large groups, orange: small groups, dark grey: lone individuals). All individual performances were averaged within the same size category (solid lines). The light-shaded areas, dark-shaded areas, and dashed curves show the 95%, 50%, and median Bayesian credible intervals of the phenomenological, time-series logistic regression. (Bottom row): Change in the main effect of the dummy variable of grouping on the decision accuracy at the phenomenological regression model. The shaded areas are the Bayesian CIs and solid curves are the median.

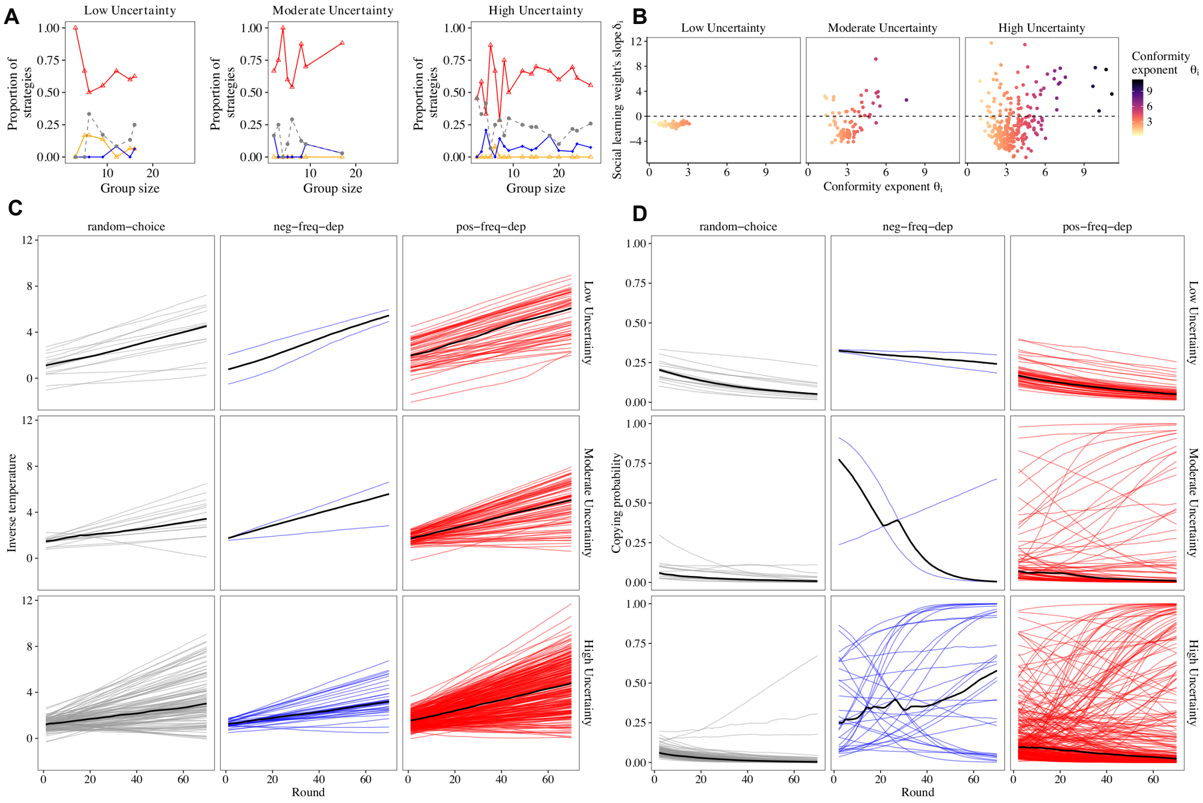


**Figure 5. Model fitting for the three different tasks uncertainty conditions**. A: Frequencies of four different learning strategies are shown in different styles (Note: orange-triangle line indicates the weak-positive frequency-dependent strategy $0 < \theta_i \leq 1$). B: The relationship between $\theta_i$ and the slope of social learning weight $\delta_i$. The horizontal dashed lines indicate a threshold at which the social learning weight $\sigma_{i,t}$ does not change with time (i.e. $\delta_i = 0$). C: Individual inverse temperature $\beta_{i,t}$ fit for each experimental participant. D: Individual social learning weight $\sigma_{i,t}$ fit for each experimental participant.

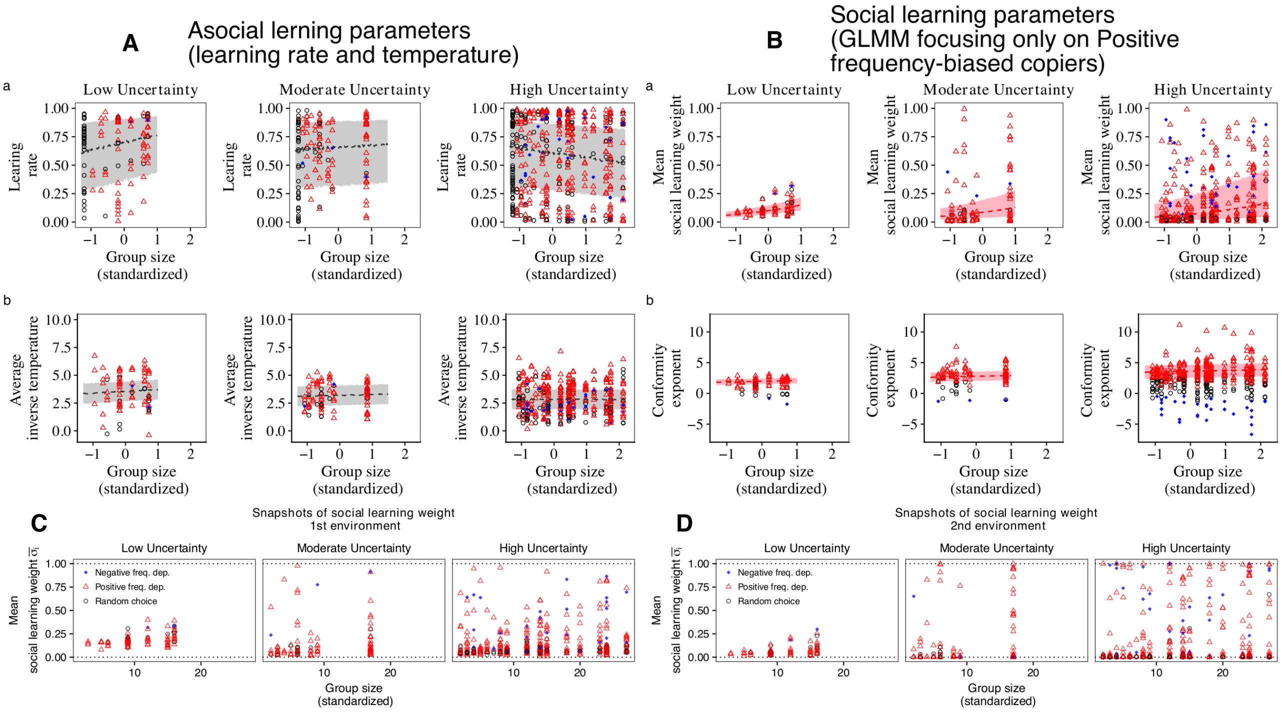


**Figure 6. Relationships between group size, task uncertainty, and fitted parameters**. A: Estimated learning rate $\alpha_i$ and estimated mean inverse temperature $\bar{\beta}_i = (\sum_t \beta_{i,t})/70$ for each individual shown for each different learning strategy (Colours are the same as in Figure 4 in the main text). The fitted generalised mixed models are shown by dashed lines (the shaded areas indicate 50% Bayesian credible intervals). B: The same figure as Figure 4B and 4C, except that the generalised mixed models were fitted only to the positive frequency-biased choice individuals (the shaded areas indicate 50% Bayesian credible intervals). C: Mean social learning weights averaged over the 1st environment ($1 \leq t \leq 40$) and D: the 2nd environment $41 \leq t$.

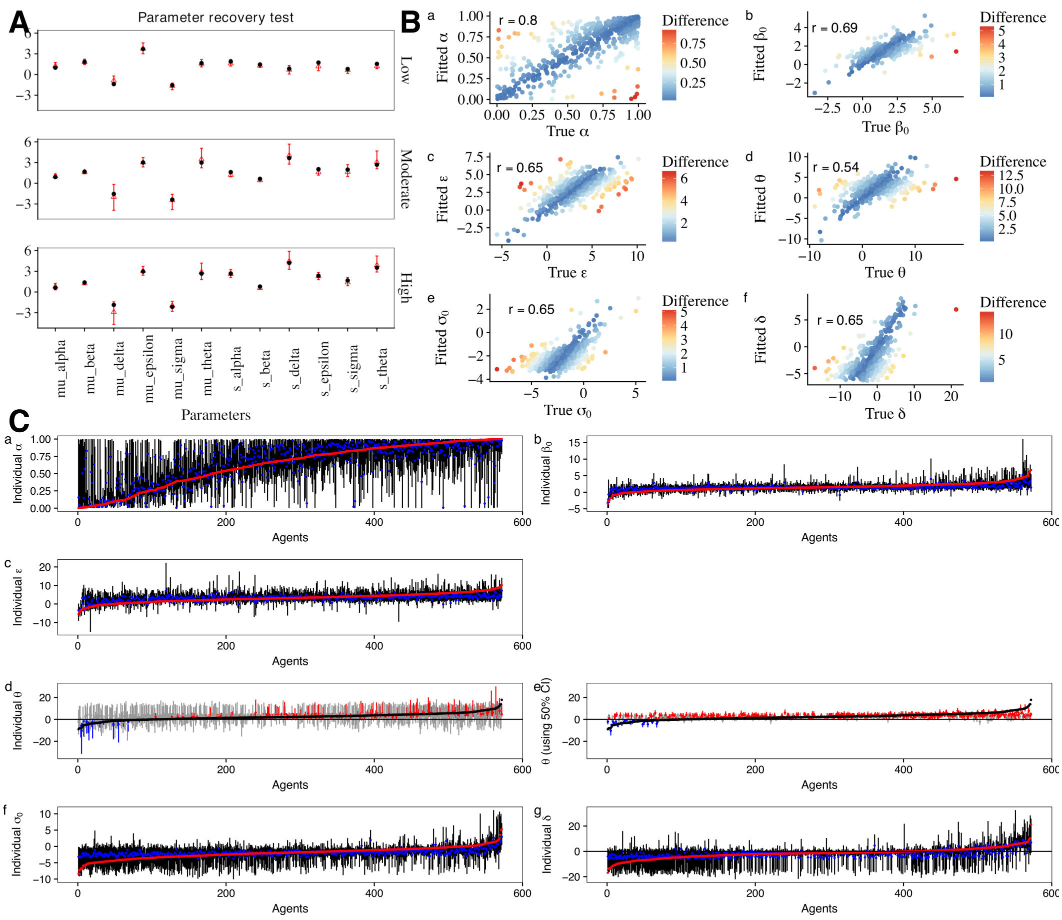


**Figure 7. The parameter recovery performance on the (A) global parameters and (B,C) individual-level parameters**. (A) The black points are the true values and the red triangles are the mean posterior values (i.e. recovered values). The 95% Bayesian credible intervals are shown by the error bars. (B) The x-axis is the true value and the y-axis is the fitted (i.e. the mean posterior) individual value. The differences between the true value and the fitted value are shown in different colours (Blue: fit well). The correlation coefficients between the true and fitted values are shown. (C) The red middle points are the true individual parameter values, the blue points are the mean posterior fitted values and the black lines are the 95% Bayesian CI. For $\theta_i$ (d & e), the true values are in black and categorisation of three different strategies, based on the 95% (d) and 50% (e) Bayesian CI, are shown in different colours (Red: positive frequency-dependent, Blue: negative frequency-dependent, Grey: asocial random decision-making.

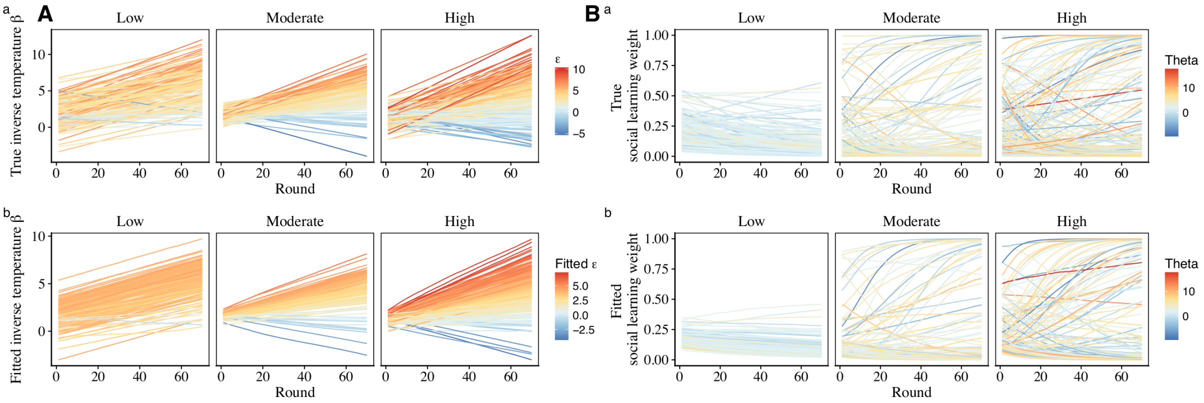

**Figure 8. The temporal evolution of (A) the individual inverse temperature $\beta_{i,t}$ and (B) social learning weight** $sigma_{i,t}$. The true parameters are shown at the top and the recovered values are shown at the bottom. The magnitude of individual slope parameter $\epsilon_i$ and conformity exponent $\theta_i$ are shown in different colours at A and B, respectively.

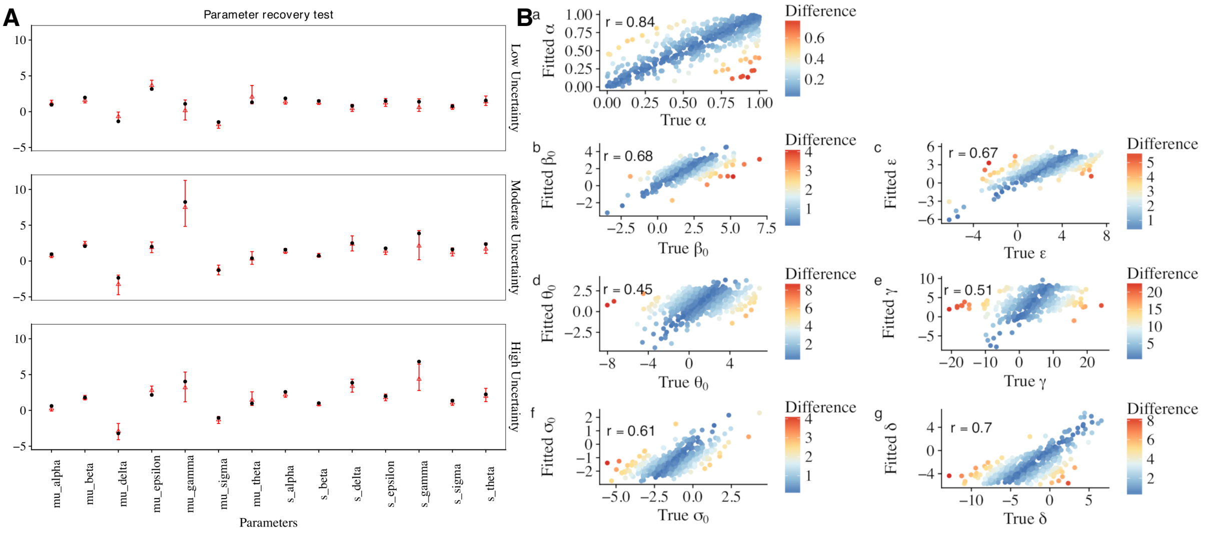

**Figure 9. The parameter recovery performance of the alternative, 'time-dependent conformity' model**. (A) The recovery of global parameters and (B) that of individual-level parameters. The presentation is the same as Figure S7 A and B.

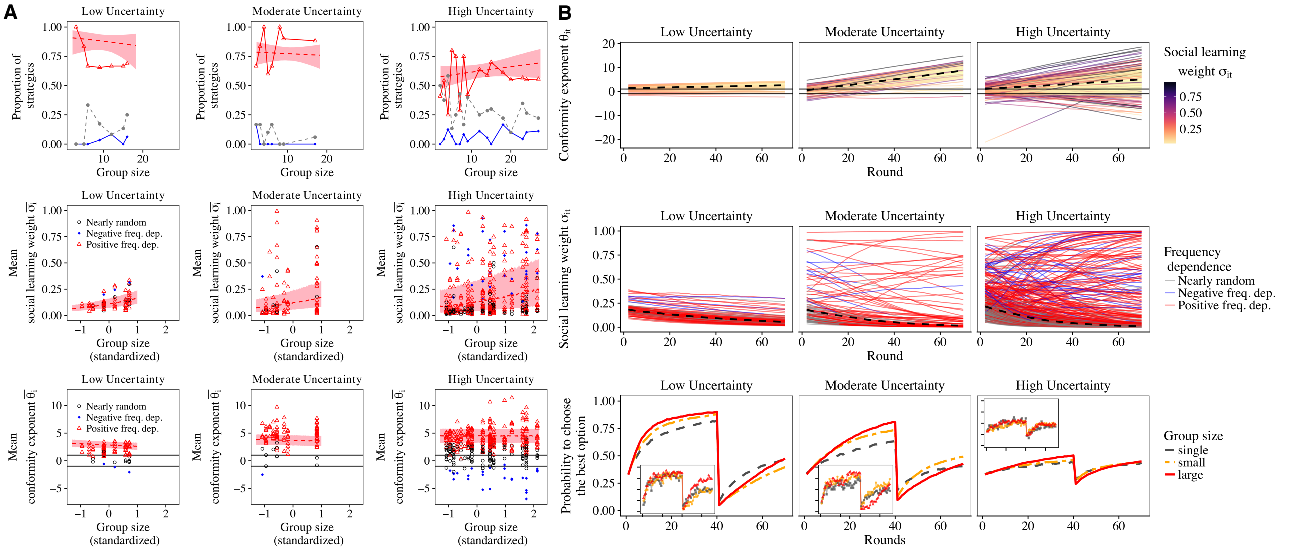


**Figure 10. Model fitting analysis using the alternative model**. (A) Model fitting for the three different task uncertainty conditions (the Low-, Moderate- and High-uncertainty) and the different group size (The same presentation as Fig. 4 in the main text). Note that the averaged conformity exponent $\bar{\theta}_i$ is used because it changes over time. (B) Change in fitted values (i.e. median of the Bayesian posterior distribution) of (Top row) the conformity exponent $\theta_{i,t}$ and (Middle row) the social learning weight $\sigma_{i,t}$ with time for each individual, for each level of task uncertainty. Thick dashed lines are the median values across the subjects for each uncertainty condition. (Bottom row) Change in average decision accuracy of the individual-based post-hoc model simulations using the experimentally fit parameter values of the alternative model (shown in main panels). The inner panels show the average decision accuracies of the experimental participants. Each line indicates different group-size categories (red-solid: large groups, orange-halfdashed: small groups, grey-dashed: lone individuals). All individual performances were averaged within the same size category (see the main text for the group sizes in each size category).

### 4 Supplementary Tables

**Supplementary Table 1**

Distributions of group sizes in the online experiment

| | Group size | | | | | | | | | | | | | | | | | | |
|---|---|---|---|---|---|---|---|---|---|---|---|---|---|---|---|---|---|---|---|
| | 1 | 2 | 3 | 4 | 5 | 6 | 7 | 8 | 9 | 12 | 14 | 15 | 16 | 17 | 18 | 20 | 23 | 24 | 27 |
| Low Uncertainty | 36 | | 1 | | 1 | 1 | | | 3 | 1 | | 1 | 1 | | | | | | |
| Moderate Uncertainty | 34 | 3 | 4 | 1 | 2 | 4 | | 1 | 1 | | | | | 2 | | | | | |
| High Uncertainty | 56 | 10 | 7 | 6 | 3 | 3 | 1 | 3 | 4 | 3 | 4 | 2 | | | 1 | 1 | 1 | 2 | 1 |

Numbers of sessions are shown for each different group size.

**Supplementary Table 2**

The mean and the 95% Bayesian credible intervals of the posterior for the group size effect at the phenomenological Gaussian model

| | Low Uncertainty | | Moderate Uncertainty | | High Uncertainty | |
|---|---|---|---|---|---|---|
| $\gamma_1$ | 0.01 | [-0.05, 0.07] | 0.02 | [-0.01, 0.05] | 0.01 | [0.00, 0.02] |
| $\gamma_2$ | 0.14 | [0.08, 0.21] | -0.05 | [-0.08, -0.01] | 0.00 | [-0.02, 0.01] |

Note: All $\hat{R}$ values are 1.0 and the effective sample sizes are larger than 883.

**Supplementary Table 3**

The mean and the 95% Bayesian credible intervals of the posterior global variance parameters governing the magnitude of individual variations in each free parameter. The number of participants ($N$) for each experimental condition are also shown.

| | Group condition | | | Solitary condition | | |
|---|---|---|---|---|---|---|
| | Uncertainty | | | Uncertainty | | |
| Parameters | Low | Moderate | High | Low | Moderate | High |
| $\nu_{\alpha^*}$ (learning rate) | 1.88 | 1.61 | 2.69 | 1.79 | 2.37 | 2.34 |
| | [1.20, 2.82] | [1.14, 2.23] | [2.23, 3.21] | [0.98, 3.03] | [1.46, 3.58] | [1.39, 3.77] |
| $\nu_{\beta_0^*}$ (inv. temp.) | 1.45 | 0.64 | 0.79 | 0.77 | 0.96 | 0.71 |
| | [0.91, 2.16] | [0.34, 1.00] | [0.61, 1.00] | [0.42, 1.25] | [0.46, 1.61] | [0.40, 1.08] |
| $\nu_{\epsilon}$ (inv. temp.) | 1.73 | 2.04 | 2.36 | 1.75 | 1.76 | 2.23 |
| | [0.41, 3.34] | [1.27, 2.96] | [1.89, 2.85] | [0.85, 2.94] | [0.85, 2.87] | [1.43, 3.08] |
| $\nu_{\sigma_0^*}$ (soc. wight) | 0.79 | 1.98 | 1.67 | – | – | – |
| | [0.17, 1.50] | [1.17, 3.15] | [1.26, 2.18] | – | – | – |
| $\nu_{\delta}$ (soc. wight) | 0.84 | 3.66 | 4.22 | – | – | – |
| | [0.05, 1.94] | [1.92, 5.78] | [3.30, 5.40] | – | – | – |
| $\nu_{\theta}$ (conformity coeff.) | 1.54 | 2.69 | 3.53 | – | – | – |
| | [0.87, 2.61] | [1.60, 4.32] | [2.67, 4.65] | – | – | – |
| $N$ | 77 | 98 | 398 | 36 | 34 | 56 |

**Supplementary Table 4**

Mean and the 95% Bayesian credible intervals of the fixed effects in the GLMM predicting the probability to become a positive frequency-dependent copier. The sized effects whose CI are either below or above zero (i.e. significant) are shown in bold face.

| Effect | 2.5% | 50% | 97.5% | Effective sample size | Rhat |
|---|---|---|---|---|---|
| $\beta_1$ **(intercept)** | 1.05 | 1.71 | 2.50 | 667 | 1.01 |
| $\beta_2$ (group size) | -0.94 | -0.05 | 0.87 | 2744 | 1.00 |
| $\beta_3$ **(uncertainty)** | -1.88 | -1.02 | -0.25 | 1548 | 1.00 |
| $\beta_4$ (age) | -0.12 | 0.43 | 1.10 | 925 | 1.01 |
| $\beta_5$ (gender) | -1.06 | -0.13 | 0.84 | 3154 | 1.00 |
| $\beta_6$ (size*uncrtn) | -0.72 | 0.24 | 1.19 | 2880 | 1.00 |
| $\beta_7$ (size*age) | -0.16 | 0.08 | 0.32 | 1869 | 1.01 |
| $\beta_8$ (size*gndr) | -0.37 | 0.03 | 0.44 | 4875 | 1.00 |
| $\beta_9$ **(uncrtn*age)** | -1.46 | -0.73 | -0.15 | 3167 | 1.00 |
| $\beta_{10}$ (uncrtn*gndr) | -1.10 | 0.02 | 1.09 | 3661 | 1.00 |
| $\beta_{11}$ (age*gndr) | -0.39 | -0.02 | 0.37 | 2712 | 1.00 |

**Supplementary Table 5**

Mean and the 95% Bayesian credible intervals of the fixed effects in the GLMM predicting individual parameter values of the social learning weight $\bar{\sigma}_i$. The sized effects whose CI are either below or above zero (i.e. significant) are shown in bold face.

| Effect | 2.5% | 50% | 97.5% | nEff | Rhat |
|---|---|---|---|---|---|
| $\beta_1$ **(intercept)** | -2.32 | -2.09 | -1.84 | 4959 | 1.00 |
| $\beta_2$ **(group size)** | 0.15 | 0.52 | 0.93 | 5230 | 1.00 |
| $\beta_3$ **(uncertainty)** | -0.98 | -0.59 | -0.22 | 4784 | 1.00 |
| $\beta_4$ **(age)** | -0.36 | -0.18 | -0.02 | 2126 | 1.00 |
| $\beta_5$ (gender) | -0.45 | -0.16 | 0.13 | 4513 | 1.00 |
| $\beta_6$ (size*uncrtn) | -0.57 | -0.10 | 0.34 | 5440 | 1.00 |
| $\beta_7$ (size*age) | -0.19 | -0.02 | 0.14 | 5359 | 1.00 |
| $\beta_8$ (size*gndr) | -0.32 | -0.01 | 0.30 | 4127 | 1.00 |
| $\beta_9$ (uncrtn*age) | -0.17 | 0.07 | 0.32 | 4088 | 1.00 |
| $\beta_{10}$ (uncrtn*gndr) | -0.37 | 0.12 | 0.62 | 4205 | 1.00 |
| $\beta_{11}$ (age*gndr) | -0.09 | 0.12 | 0.35 | 4963 | 1.00 |
| $\gamma$ **(uncertainty effct on variance)** | 1.11 | 1.38 | 1.62 | 3067 | 1.00 |

**Supplementary Table 6**

Mean and the 95% Bayesian credible intervals of the fixed effects in the GLMM predicting the social learning weight $\bar{\sigma}_i$ for the positive frequency-biased choice individuals only. The sized effects whose CI are either below or above zero (i.e. significant) are shown in bold face.

| Effect | 2.5% | 50% | 97.5% | nEff | Rhat |
|---|---|---|---|---|---|
| $\beta_1$ **(intercept)** | -2.42 | -2.17 | -1.91 | 5601 | 1.00 |
| $\beta_2$ **(group size)** | 0.09 | 0.47 | 0.90 | 4509 | 1.00 |
| $\beta_3$ (uncertainty) | -0.75 | -0.28 | 0.17 | 6011 | 1.00 |
| $\beta_4$ (age) | -0.33 | -0.14 | 0.04 | 5796 | 1.00 |
| $\beta_5$ (gender) | -0.36 | -0.03 | 0.30 | 6075 | 1.00 |
| $\beta_6$ (size*uncrtn) | -0.55 | -0.01 | 0.49 | 5410 | 1.00 |
| $\beta_7$ (size*age) | -0.27 | -0.06 | 0.14 | 6022 | 1.00 |
| $\beta_8$ (size*gndr) | -0.42 | -0.05 | 0.33 | 6174 | 1.00 |
| $\beta_9$ (uncrtn*age) | -0.36 | -0.04 | 0.29 | 6483 | 1.00 |
| $\beta_{10}$ (uncrtn*gndr) | -0.75 | -0.13 | 0.49 | 4746 | 1.00 |
| $\beta_{11}$ (age*gndr) | -0.16 | 0.10 | 0.36 | 5927 | 1.00 |
| $\gamma$ **(uncertainty effct on variance)** | 1.14 | 1.50 | 1.80 | 5729 | 1.00 |

**Supplementary Table 7**

Mean and the 95% Bayesian credible intervals of the fixed effects in the GLMM predicting individual parameter values of the conformity exponent $\theta_i$. The sized effects whose CI are either below or above zero (i.e. significant) are shown in bold face.

| Effect | 2.5% | 50% | 97.5% | nEff | Rhat |
|---|---|---|---|---|---|
| **$\beta_1$ (intercept)** | 1.30 | 1.64 | 2.01 | 2571 | 1.00 |
| $\beta_2$ (group size) | -0.69 | -0.17 | 0.35 | 5443 | 1.00 |
| **$\beta_3$ (uncertainty)** | 0.38 | 0.90 | 1.41 | 2602 | 1.00 |
| $\beta_4$ (age) | -0.19 | 0.07 | 0.33 | 2967 | 1.00 |
| $\beta_5$ (gender) | -0.34 | 0.10 | 0.54 | 3557 | 1.00 |
| $\beta_6$ (size*uncrtn) | -0.40 | 0.19 | 0.79 | 5317 | 1.00 |
| $\beta_7$ (size*age) | -0.27 | -0.06 | 0.14 | 5172 | 1.00 |
| $\beta_8$ (size*gndr) | -0.24 | 0.13 | 0.50 | 5167 | 1.00 |
| $\beta_9$ (uncrtn*age) | -0.59 | -0.26 | 0.07 | 3436 | 1.00 |
| $\beta_{10}$ (uncrtn*gndr) | -0.86 | -0.21 | 0.45 | 3509 | 1.00 |
| $\beta_{11}$ (age*gndr) | -0.30 | -0.02 | 0.27 | 4885 | 1.00 |
| **$\gamma$ (uncertainty effct on variance)** | 1.07 | 1.31 | 1.54 | 4178 | 1.00 |

**Supplementary Table 8**

Mean and the 95% Bayesian credible intervals of the fixed effects in the GLMM predicting the conformity exponent $\theta_i$ for the positive frequency-biased choice individuals only. The sized effects whose CI are either below or above zero (i.e. significant) are shown in bold face.

| Effect | 2.5% | 50% | 97.5% | nEff | Rhat |
|---|---|---|---|---|---|
| $\beta_1$ **(intercept)** | 1.74 | 2.00 | 2.29 | 4922 | 1.00 |
| $\beta_2$ (group size) | -0.40 | 0.03 | 0.42 | 5695 | 1.00 |
| $\beta_3$ **(uncertainty)** | 1.20 | 1.64 | 1.04 | 4381 | 1.00 |
| $\beta_4$ (age) | -0.32 | -0.13 | 0.05 | 6046 | 1.00 |
| $\beta_5$ (gender) | -0.40 | -0.07 | 0.26 | 5988 | 1.00 |
| $\beta_6$ (size*uncrtn) | -0.47 | 0.00 | 0.50 | 4458 | 1.00 |
| $\beta_7$ (size*age) | -0.24 | -0.07 | 0.11 | 5716 | 1.00 |
| $\beta_8$ (size*gndr) | -0.12 | 0.19 | 0.51 | 5349 | 1.00 |
| $\beta_9$ (uncrtn*age) | -0.15 | 0.10 | 0.37 | 6424 | 1.00 |
| $\beta_{10}$ (uncrtn*gndr) | -0.53 | -0.01 | 0.51 | 5710 | 1.00 |
| $\beta_{11}$ (age*gndr) | -0.14 | 0.09 | 0.33 | 6545 | 1.00 |
| $\gamma$ **(uncertainty effct on variance)** | 0.71 | 0.91 | 1.10 | 6545 | 1.00 |

**Supplementary Table 9**

Mean and the 95% Bayesian credible intervals of the fixed effects in the GLMM predicting individual parameter values of the learning rate $\alpha_i$. The sized effects whose CI are either below or above zero (i.e. significant) are shown in bold face.

| Effect | 2.5% | 50% | 97.5% | nEff | Rhat |
|---|---|---|---|---|---|
| **$\beta_1$ (intercept)** | 0.12 | 0.67 | 1.23 | 4887 | 1.00 |
| $\beta_2$ (group size) | -0.49 | 0.24 | 0.96 | 5413 | 1.00 |
| $\beta_3$ (uncertainty) | -0.93 | -0.27 | 0.38 | 4827 | 1.00 |
| $\beta_4$ (age) | -0.48 | -0.03 | 0.40 | 5794 | 1.00 |
| $\beta_5$ (gender) | -0.40 | 0.38 | 1.17 | 4908 | 1.00 |
| $\beta_6$ (size*uncrtn) | -1.24 | -0.48 | 0.29 | 5423 | 1.00 |
| $\beta_7$ (size*age) | -0.25 | -0.04 | 0.17 | 6157 | 1.00 |
| $\beta_8$ (size*gndr) | -0.25 | 0.15 | 0.54 | 6305 | 1.00 |
| $\beta_9$ (uncrtn*age) | -0.09 | 0.38 | 0.85 | 6085 | 1.00 |
| $\beta_{10}$ (uncrtn*gndr) | -1.21 | -0.29 | 0.65 | 4699 | 1.00 |
| $\beta_{11}$ (age*gndr) | -0.35 | -0.01 | 0.34 | 5824 | 1.00 |

**Supplementary Table 10**

Mean and the 95% Bayesian credible intervals of the fixed effects in the GLMM predicting individual parameter values of the average inverse temperature $\bar{\beta}_i$. The sized effects whose CI are either below or above zero (i.e. significant) are shown in bold face.

| Effect | 2.5% | 50% | 97.5% | nEff | Rhat |
|---|---|---|---|---|---|
| $\beta_1$ **(intercept)** | 3.09 | 3.47 | 3.85 | 5906 | 1.00 |
| $\beta_2$ (group size) | -0.48 | 0.03 | 0.54 | 5707 | 1.00 |
| $\beta_3$ (uncertainty) | -0.87 | -0.43 | 0.02 | 5863 | 1.00 |
| $\beta_4$ (age) | -0.49 | -0.21 | 0.08 | 4498 | 1.00 |
| $\beta_5$ (gender) | -0.35 | 0.16 | 0.67 | 5845 | 1.00 |
| $\beta_6$ (size*uncrtn) | -0.73 | -0.17 | 0.37 | 5503 | 1.00 |
| $\beta_7$ (size*age) | -0.20 | -0.06 | 0.08 | 6454 | 1.00 |
| $\beta_8$ **(size*gndr)** | 0.02 | 0.26 | 0.50 | 6492 | 1.00 |
| $\beta_9$ **(uncrtn*age)** | 0.01 | 0.33 | 0.63 | 6167 | 1.00 |
| $\beta_{10}$ (uncrtn*gndr) | -1.19 | -0.58 | 0.02 | 5718 | 1.00 |
| $\beta_{11}$ (age*gndr) | -0.33 | -0.10 | 0.12 | 5558 | 1.00 |
